# GCS-H2 is essential for growth as it acts as the main relay for mitochondrial lipoylation in heterotrophic tissues of *Arabidopsis thaliana*

**DOI:** 10.1101/2024.10.14.617781

**Authors:** Jonathan Przybyla-Toscano, Clément Boussardon, Marta Juvany, Saleh Alseekh, Katharina Modde, Aurélien Valentin, Hans Stenlund, Hermann Bauwe, Alisdair R Fernie, Olivier Keech

**Author notes:** Univ. Grenoble Alpes, CEA, INRAE, CNRS, IRIG, LPCV, 38000 Grenoble, France. Equally contributed (Jonathan Przybyla-Toscano, Clément Boussardon).

## Abstract

H-protein is part of the multi-protein glycine cleavage system found in both prokaryotes and eukaryotes. The *Arabidopsis thaliana* genome contains three loci for genes encoding H-protein. We show that in land plants, there are two clades of mitochondrial H-proteins; one that clusters with H1 and H3, and another that clusters with H2, an isoform mostly expressed in heterotrophic tissue and for which a null mutation results in a >99% impeded growth. After showing that all three isoforms fulfill the same metabolic function, we evidenced that the impaired root growth in *h2* results from an altered cell cycle progression concomitant to a lower Target Of Rapamycin kinase activity. Subsequent metabolomic approaches revealed an accumulation of storage sugars and a significant decrease of the TCA intermediates and several vitamins in the *h2* root cells as compared to the wild-type ones.

Additional investigations evidenced that H2 acts as the main relay for lipoylation in root mitochondria, which diverges from the current lipoylation model proposed for photosynthetic tissues. Together, this work provides new insight on the control of the cell cycle by mitochondrial metabolism, and also challenges the current dogma for lipoylation in mitochondria of plant cells.

**Once sentence summary:** The absence of GCS-H2, an isoform of H-protein found in heterotrophic tissue, impedes the cell cycle, and drastically affects plant growth as it acts as the main relay for mitochondrial lipoylation.

## INTRODUCTION

The glycine cleavage system (GCS), also known in plants as the glycine decarboxylase complex, is found in most of organisms, both prokaryotes and eukaryotes. This complex is involved notably in the formation of one-carbon (C1) units, which are essential for nucleotide synthesis, amino acid homeostasis, epigenetic maintenance and reductive metabolism (Hanson and Roje, 2001; Ducker and Rabinowitz, 2017; Mutka et al 2022). GCS comprises four proteins including three enzymes designated as P-protein (a glycine decarboxylase *per se*), T-protein (an aminomethyl-transferase), L-protein (a dihydrolipoamide dehydrogenase) and a carrier lipoic acid-dependent protein named H-protein. These multi-step reactions have been discussed in detail for several species (Kikuchi and Hiraga 1982; Oliver 1994; Douce et al 2001).

In the photosynthetic cells of plants, GCS is present in high abundance as it also contributes to the photorespiratory cycle, a complex pathway that detoxifies and recycles the phosphoglycolate resulting from the oxygenase activity of Ribulose 1,5 Bisphosphate Carboxylase/Oxygenase (RUBISCO) (Oliver et al 1990; Bauwe et al 2010). Biochemically speaking, GCS catalyses the tetrahydrofolate (THF)-dependent oxidation and deamination of glycine leading to the release of CO^2^, NH^3^ and a concomitant reduction of NAD^+^ to NADH. The remaining carbon of glycine is transferred to THF, and forms 5,10-methylene-THF (also defined as CH^2^-THF). In the case of photorespiratory metabolism, CH^2^-THF reacts with another glycine to form serine in a reversible reaction catalysed by serine hydroxymethyltransferase (SHMT).

In C^3^ plants, it was primarily reported that GCS mutants were lethal only under photorespiratory conditions, and no differences to the wild type (WT) were observed when these mutants were grown under elevated CO^2^; a condition that suppresses the oxygenase activity of RUBISCO and, in turn, glycine synthesis from phosphoglycolate (Sommerville and Ogren 1982; Blackwell et al 1990). With advanced genome identification, it became evident that several isoforms of GCS subunits were present in plants, and that this functional redundancy contributed to the non-lethality. For instance, in 2007, Engel et al. demonstrated that while individual knockouts of the two genes coding for P-protein isoforms were barely affected in their development, the double mutant did not develop beyond the cotyledon stage in air enriched with 0.9% CO^2^, and died after a few weeks. Despite this interesting observation not all isoforms of GCS subunits have been studied to date.

The genome of Arabidopsis contains three loci for genes encoding H-protein: H1 – *At1g32470*, H2 – *At2g35120* and H3 – *At2g35370*. According to the expression data of the AtGenExpress Consortium (Arabidopsis eFP Browser, http://bar.utoronto.ca/; SFig. 1), the genes coding for *H1* and *H3* are principally expressed in photosynthetic tissues, while *H2* is mostly expressed in heterotrophic tissues i.e. dry seeds and roots. In line with these expression data, only the H2 isoform was detected in isolated mitochondria from a heterotrophic Arabidopsis cell suspension culture (Fuchs et al 2020). The aim of the present work was to delineate the role of the H2 isoform in Arabidopsis plants. We successfully isolated two independent homozygous T-DNA insertion mutants, which exhibit a drastic growth alteration, even under non-photorespiratory conditions. Although the three H isoforms differ in their amino sequence, both constitutive and promoter-specific expression of *H1* and *H3* can fully rescue the dwarf phenotype of the *h2* mutant, thus demonstrating that a similar metabolic function is shared between the three isoforms. In a second step, we showed that the strikingly impeded root growth observed in *h2* results from an altered cell cycle progression as evidenced by a lower Target Of Rapamycin (TOR) kinase activity in root cells. Subsequent metabolomic approaches revealed major differences in the *h2* root cells as compared to the wild-type (WT) ones. In particular, a significant decrease of both TCA intermediates and vitamins was observed, whereas storage sugars, glycine and branched chain amino acids (BCAA) accumulated. While several metabolic hypotheses were progressively tested and ruled out, we discovered that the lipoylation of the E2 subunits of 2-oxoacid dehydrogenase complexes was severely affected in the mutant, which demonstrates, in direct contrast to the current model for lipoylation in mitochondria of photosynthetic cells, that H2 acts as the main relay for mitochondrial lipoylation in root cells.

## RESULTS

### GCS-Hs are evolutionarily conserved proteins

As stated above, the Arabidopsis genome contains three independent loci coding for H-proteins: *At1g32470* for H1; *At2g35120* for H2 and *At2g35370* for H3. Alignment of Arabidopsis H-protein primary sequences with bacterial, yeast and human orthologs reveal a strict conservation of residues essential for their function i.e. the conserved lysine required for lipoamide binding as well as hydrophobic residues making contacts with the lipoamide chain (Cohen-Addad et al 1995; Fig. 1a). Interestingly, mature H1 and H3 proteins from Arabidopsis have a very high (95%) sequence identity, whereas mature H2 is atypical since it shares only 67% and 69% identity with H1 and H3, respectively (SFig. 2). A phylogenetic analysis revealed that H-proteins are evolutionarily conserved from algae to land plants reflecting important functions for this protein (Fig. 1b). While green algae possess a single H-protein gene, most land plant species acquired additional *H* genes. On the basis of protein sequences, a phylogenetic tree showed that H-proteins from higher plants are grouped into two clades in which H2 was clearly distinguishable from the H1/H3 homologs (Fig. 1b).

**Figure 1.**
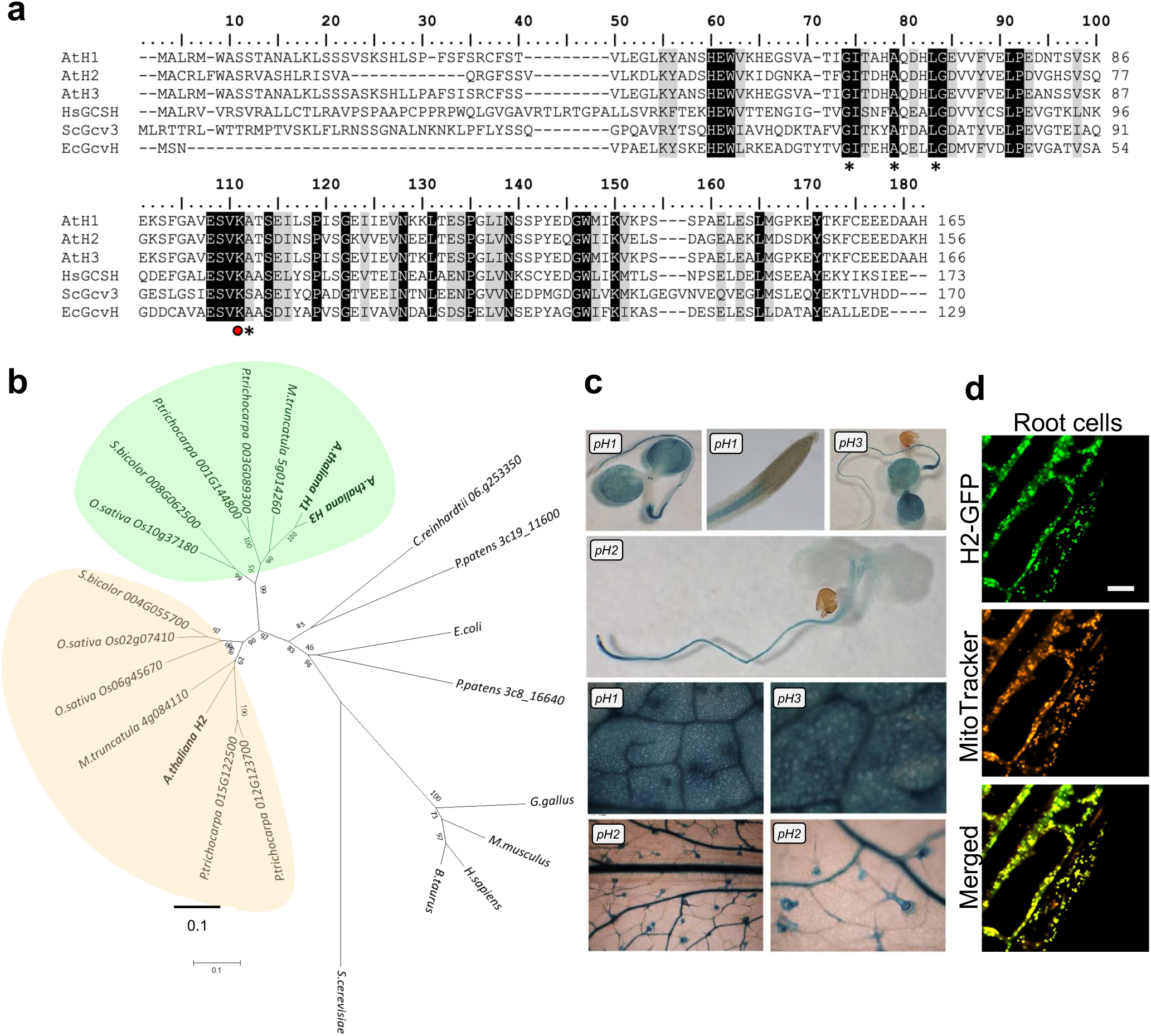
Amino acid sequence alignment and phylogeny of H proteins. (**a**) Alignment was obtained by using ClustalW (BioEdit). *A. thaliana* (At), *H. sapiens* (Hs), *E. coli* (Ec) and *S. cerevisiae* (Sc). Conserved lipoyl-lysine is indicated by a red dot, whereas hydrophobic residues making contacts with the lipoamide chain (Cohen-Added et al., 1995) are indicated by an asterisk. **(b)** Phylogenetic analysis of H family. Evolutionary analyses were conducted in MEGA7. The tree was constructed using the Neighbor-Joining method. Branch lengths are proportional to phylogenetic distances. The percentage of replicate trees in which the associated taxa clustered together in the bootstrap test (1000 replicates) are shown next to the branches.(**c**) GUS assays with *pH1-, pH2- and pH3::GUS* showing the tissue-specific expression of genes coding H-proteins in Arabidopsis. (**d)** Subcellular localization of H2. Expression of *pH2::H2:GFP* in root cells from a 14-day-old seedling. Scale bar: 10 µm.

### H2 is a mitochondrial protein that is most abundant in heterotrophic tissues

The tissue-specific expression of the three isoforms was studied using GUS reporter constructs (Fig. 1c). At an early development stage, i.e. in 7-day-old transgenic seedlings, the *H2* promoter had an activity restricted to roots while promoters of *H1 and H3* showed expression in cotyledons and young roots, although not in the root cap (Fig. 1c). In mature leaves, GUS activity was detected in vascular bundles and trichomes only in the case of *H2,* while a more diffuse signal was observed from both vascular bundles and mesophyll cells in the case of *H1* and *H3*. These results corroborate the expression levels retrieved from the AtGenExpress Consortium (SFig. 1), and confirmed that H-proteins are differentially expressed in Arabidopsis tissues, with *H2* being mostly expressed in heterotrophic tissues. Even though H-proteins from various eukaryotes, including plants and mammals, were previously detected in mitochondria (Fujiwara et al 1979; Oliver et al 1990; Pares et al 1994), the subcellular localization of H2 has never been specifically investigated, and according to N-terminome a shorter mitochondrial targeting sequence data is predicted for this isoform in comparison to H1 and H3 (Carrie et al 2015; Venne et al 2015). A translational fusion between the full genomic region corresponding to the *H2* gene including its cis-regulatory sequence (-1000 bp upstream of the translational start) in frame with the *GFP* reporter gene showed a fluorescent signal exhibiting a punctate pattern which mostly co-localized with a MitoTracker dye was detectable in root cells from transgenic Arabidopsis 14-day-old seedlings (Fig. 1d), hence confirming the mitochondrial localization of H2.

### *h2* loss-of-function mutants have strongly impaired growth

To determine the physiological role of H2, we isolated two mutant alleles with a T-DNA insertion in the 5’UTR region for *h2-1* or in the first exon for *h2-2* (Alonso et al 2003; Fig. 2a). When grown on soil at ambient air (i.e. 400 ppm CO^2^) and under long-day conditions, the two homozygous mutant lines carrying a recessive mutation showed a drastic growth reduction of aerial part compared to WT plants, even though the dwarf phenotype was more severe for *h2- 2* (Fig. 2b). After 7 weeks of growth, the shoot biomass of *h2-1* reached less than 1% of that of WT, a trend that was not rescued when plants were grown under high CO^2^ (2000 ppm), a condition that alleviates photorespiration (SFig. 3). Both *h2* mutant plants presented a purple pigmentation reflecting an accumulation of anthocyanin (Fig. 2b) as well as very small siliques, which led to a massively reduced seed production in *h2-1* and to a complete sterility in *h2-2*. Although the exacerbated phenotype of *h2-2* suggested that *h2-1* may be a leaky allele, which potentially has a residual level of H2 protein, neither RT-PCR nor immunoblot analyses could evidence the presence of transcript or protein, respectively, in the shoots and roots of this mutant allele (Fig. 2c-d). Indeed, the immunoblot analysis using polyclonal anti-H purified IgG showed that the upper immune signal (17-18 kDa) was systematically absent in shoot and root samples of the *h2-1* mutant. However, this was not the case for the lower signal (14-15 kDa) that remained detectable in shoot samples from WT and *h2-1* mutant (Fig. 2d). Thus, the upper band, ca. 17-18 kDa, was assigned to the native H2 while the immune signal detected at ca. 15 kDa corresponded to a mixture of H1 and H3. This different gel mobility was unexpected given that the determination of cleavage sites from Arabidopsis mitochondrial peptidases (Carrie et al 2015) suggests that all three isoforms have a predicted molecular mass of 14.5 kDa. That said, several large-scale proteomic studies have also reported a similar migration pattern for Arabidopsis H proteins (Lee et al. 2008; Ströher et al. 2016). It seems unlikely that the processing of the C-terminal part is the cause of this atypical migration because this region is very well conserved between the Arabidopsis H proteins (Fig. 1a). One of the reasons could be linked to the isoelectric point (pI) of the mature proteins, with H2 being slightly more acidic (with a theorical pI of 4.55) than H1 (4.73) and H3 (4.65).

**Figure 2.**
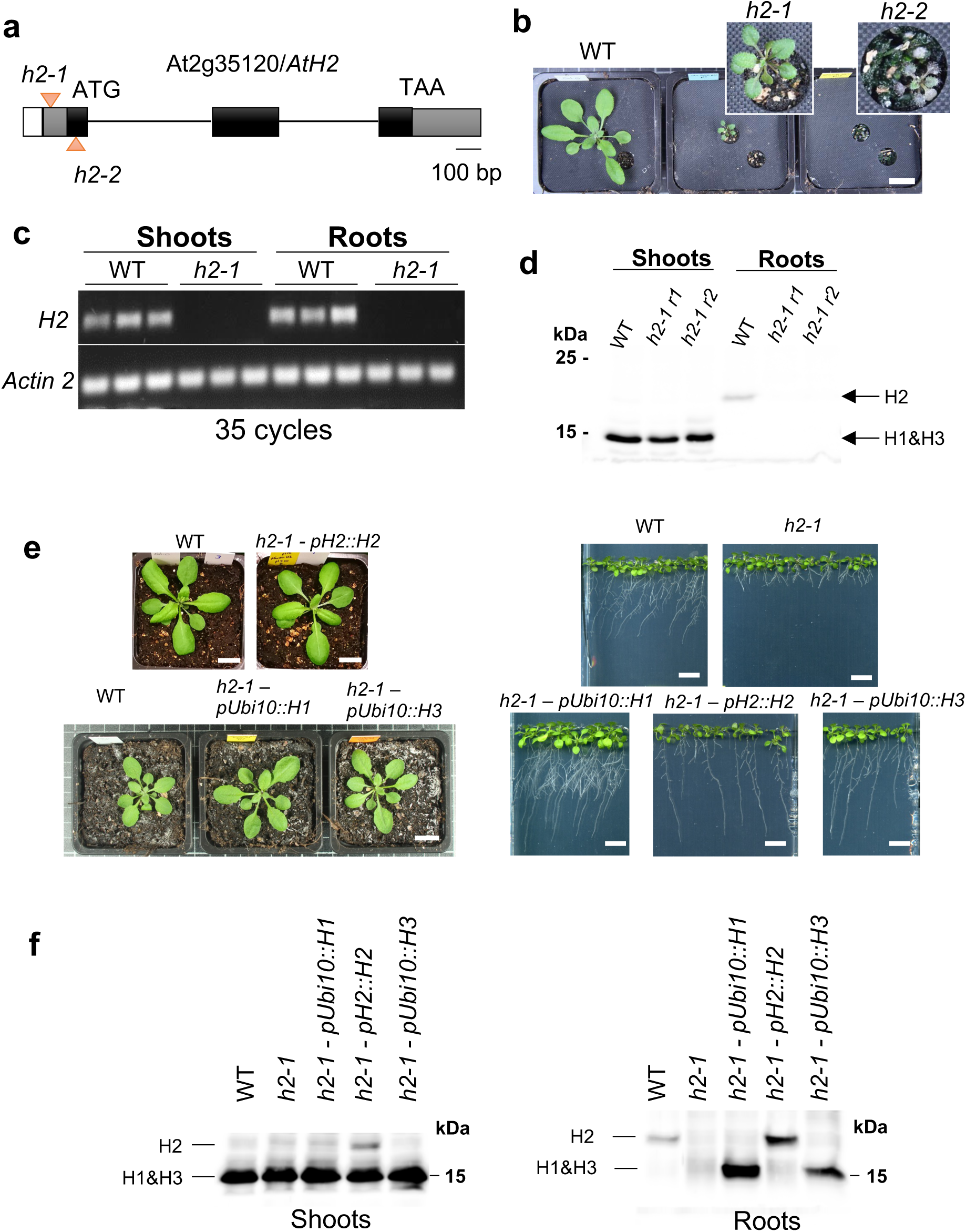
Molecular characterization of *h2* mutants from *Arabidopsis thaliana*. **(a)** Gene model of At2g35120 encoding for the H2 protein. Exons are represented in black, both 5’ and 3’ untranslated (UTRs) regions are shown in grey. Positions of T-DNA insertions of the mutant lines are represented by triangles. *h2-1* : SALKseq_120199 ; *h2-2* : SAIL_1152_G07. **(b)** Aerial growth phenotypes of 3-week-old wild-type Col-0 (WT), *h2-1* and *h2-2* plants. Scale bars = 1cm (**c**) Transcript abundance of *H2* estimated by RT-PCR after 35 cycles; *Actin 2* was used as a control. **(d)** H-proteins levels shown by immunolabelling (anti-H-protein) in a total protein extract from 2-week-old seedlings (pool of 20 seedlings) of WT and 2 independent replicates of *h2-1 (r1 and r2)*; 50 µg of soluble proteins per well. (**e**) Complementation of the *h2* mutant phenotype by the expression of H isoforms. Macro-phenotype comparison of WT and *h2-1* knockout mutant expressing i) the full-length *H2* cDNA (1000 nt upstream of the initial codon to 500 nt downstream of the stop codon) or ii) the genomic sequence of *H1* (At2g35370) or iii) of *H3* (At1g32470), both under the control the Arabidopsis ubiquitin-10 promoter (*pUbi10*). Pictures are 3-week-old-plants grown under long-day conditions; scale bar = 1 cm. (**f**) H-protein levels in WT Col- 0 and complemented *h2-1* lines used in (e).

### Ubiquitous expression of H1 or H3 isoforms rescues the *h2* phenotype

The immunoblot analysis (Fig. 2d) indicated that WT and *h2-1* had similar amount of H1+H3 in shoots while none of these two isoforms were detected in roots, which was in line with the transcript abundance data (SFig. 1). This also evidences that endogenous levels of H1 and H3 in shoot cannot compensate for the lack of H2. Since H2 shares less than 70% identity with the two other isoforms, we tested whether a functional divergence among Arabidopsis H proteins could explain the phenotype of the *h2* mutant. As the two *h2* alleles provided a strongly altered growth phenotype and *h2-2* was sterile, we elected to pursue our investigations using only the *h2-1* allele. To challenge the functional divergence hypothesis, we first generated *h2- 1* complemented lines in which *H2* expression was driven by its own promoter (1000 bp upstream of the ATG). When grown on soil or on plates, *h2-1–pH2::H2* plants appeared fully complemented with a similar rosette than the WT after 3 weeks of growth under long days conditions, and with a recovery of the root growth when placed on ½ strength MS medium (Fig. 2e). This demonstrated that the phenotypes observed in *h2-1* mutant plants were solely the result of a loss-of-function of *H2*. Secondly, *H1* or *H3* were expressed ubiquitously under the control of the constitutive promoter *ubiquitin-10* (*pUbi10*). The expression of *H1* or *H3*, respectively, allowed for the full rescue of both the dwarf rosette and the primary root growth phenotypes of the *h2-1* mutant (Fig. 2e). An immunoblot analysis using the anti-H antibodies confirmed the successful expression and translation of all the aforementioned transgenes, as well as the atypical migration pattern of H2 (Fig. 2f). Taken together, these results demonstrate that: i) H1 and H3 isoforms can fulfil similar biochemical functions as H2 protein, ii) only the spatial expression pattern of H2 is responsible of the altered growth of the *h2-1* mutant, and iii) the endogenous activity of H1 and H3 in photosynthetic tissue cannot compensate for the lack of H2 in heterotrophic tissues. The latter point implies that even though the lack of H2 could lead to a shortage of one or several key metabolite(s), the resulting metabolic imbalance cannot be rescued by an import from the photosynthetic cells.

### H2 shows a strong alteration of root growth due to cell cycle arrest

As shown in Figure 2e, a massive root growth alteration was noticed in *h2-1*. After 10 days, root growth in the *h2-1* mutant was hindered by 70% as compared to WT seedlings (Fig. 3a-b). Interestingly, etiolated seedlings of WT and *h2-1* were phenotypically undistinguishable and characterized by similar root and hypocotyl growth, which suggests that cell division, and not cell elongation, might be compromised in the *h2* mutant (Fig. 3c-d). To test this hypothesis, we transformed WT and mutant plants with a reporter construct, *pCycB1-eGFP* (Cyclin B1; Kobayashi et al 2015), that shows fluorescence if cells are able to undergo G2 and M-phases. Using confocal laser microscopy, the fluorescence was estimated in an area going from the base of the quiescent centre to 250 µm up in the root, defined as region of interest (ROI) (Fig. 3e, SFig. 4). A much lower fluorescence/ROI (p < 0.0047) was detected in the mutant, therefore confirming a compromised cell cycle. Since H2 is a mitochondrial protein and mitochondria are central organelles for the energy maintenance, we also assessed whether the activity of TOR (Target Of Rapamycin), a well-characterized kinase that regulates the cell cycle based on the energy homeostasis of the cell (Ren et al, 2012), could be affected in *h2-1*. The kinase activity of TOR, estimated by the ratio between the phosphorylated and non-phosphorylated forms of Ribosomal Protein S6 (RPS6), a major target of TOR (Dobrenel et al 2016), was not significantly different in the shoots between WT and the *h2-1* mutant. However, it was found to be significantly lower (P< 0.00027) in roots of *h2-1* as compared to the WT (Fig. 3g-h, SFig. 5).

**Figure 3.**
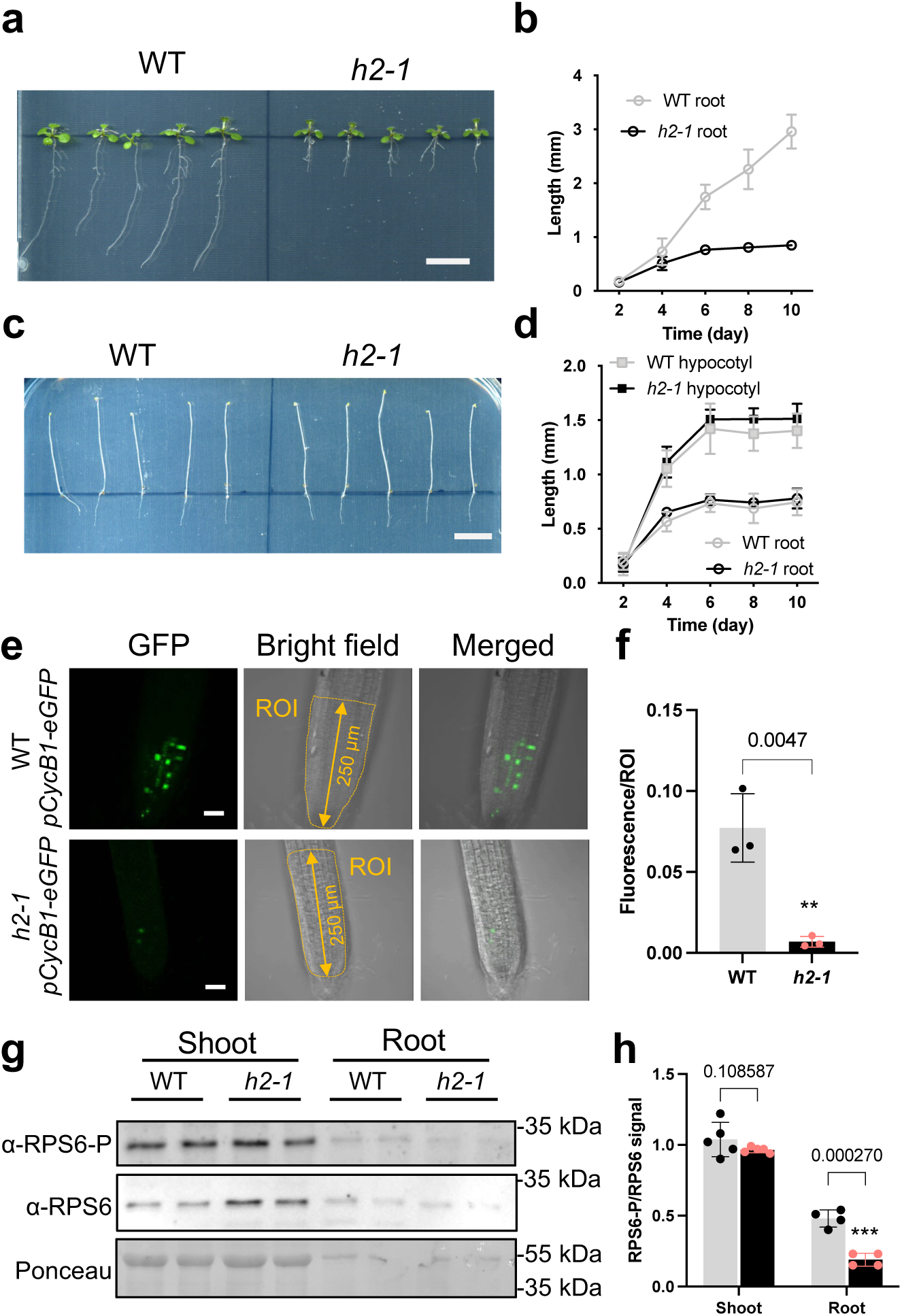
**(a)** Primary root growth in 10 day-old WT-Col0 and *h2-1* plants grown under long-day conditions; scale bars = 1cm; and (b) quantification of root growth over time for WT (Col-0) and *h2-1* plants grown in light. **(c**) Primary root and hypocotyl growth in seedlings etiolated for 10 days; scale bars = 1cm; and (**d**) quantification of root and hypocotyl growth over time for WT (Col-0) and *h2-1* plants grown in darkness. (**e**) Cell division in WT and *h2-1* root tips reported by fluorescence of GFP in root tips of 8-day-old plantlets transformed with *pCycB1- eGFP*; scale bar = 40 µm. Region of Interest (ROI) was defined as being the root area from the quiescent center to 250 µm upward as shown by the orange area defined on the bright field pictures (**f**) Quantification of fluorescence in WT and *h2-1* lines transformed with a *pCycB1-eGFP* (n=3; see SFig. 4), **: *P value* < 0.01 after Student’s t-test. (**g**) Immunoblot reporting the phosphorylation state of RPS6, target of TOR (whole gels and replicates can be found in SFig. 5). (**h**) quantification of TOR kinase activity with RPS6-P/RPS6 ratio, (n=5), ***: *P < 0.001 value* after Student’s t-test.

### *h2* mutation causes a massive metabolic shift

To elucidate how the lack of H2 in heterotrophic tissues leads to such an impaired growth, we carried out a set of complementary metabolomic analyses on both root and shoot tissues from WT and *h2-1* mutant plants. To optimize the sampling strategy (speed, amount of material and reproducibility), plants were grown hydroponically under controlled conditions at either 400 ppm or 2000 ppm CO^2^ (cf Materials and Methods). This way a potential contribution of H2 to photorespiration could be also evaluated. We first used an untargeted large spectrum GC-MS approach (Dataset S1), which showed with a subsequent Principal Component Analysis (PCA) that shoot samples did not differ much between WT and *h2-1*, regardless of the CO^2^ level applied. This ruled out any substantial role of H2 during photorespiration and in photosynthetic organs in general. In contrast, the PCA revealed that the metabolome of root cells in *h2-1* differs significantly from the one of WT. Yet, CO^2^ levels did not strongly influence this separation (Fig. 4a). An additional Orthogonal Partial Least Squares Discriminant Analysis (OPLS-DA) was used for supervised classification (Bylesjö et al 2006). The Shared and Unique Structure (SUS) plot was used for metabolite contribution comparisons between models (Fig. 4b). This revealed that, as compared to WT, root cells of *h2-1* at either 400 or 2000 ppm contained significantly lower levels of TCA intermediates and several vitamins such as pantothenic acid or ascorbate, while glycine, BCAA, pyruvate and storage sugars such as maltose, trehalose, raffinose were significantly more abundant (Fig. 4b; Dataset S1). Placing several of these metabolites in known metabolic pathways allowed a better visualization of the metabolic shift that occurs in *h2-1* root cells, as well as to raise a few hypotheses to clarify the metabolic events that lead to impaired growth in *h2-1* (Fig. 4c). Based on our results, we primarily raised three hypotheses (Fig. 4c):

1. Due to metabolic constrains, the lower TOR kinase activity could result from its repression by SNF1-related protein kinase (SnRK1) via a varying level of trehalose-6- phosphate (T6P), in a loop where T6P represses SnRK1, which in turn represses TOR (Figueroa and Lunn, 2016; Baena-González and Hanson, 2017). In such model (Fig. 4c), one would therefore expect that the level of T6P is lower in *h2-1* than in WT root cells;
2. CH^2^-THF, which is a direct product from the decarboxylation of glycine via GCS is necessary for the biosynthesis of vitamin B5, also known as pantothenic acid. Pantothenic acid is notably required for the biosynthesis of coenzyme-A (CoA). Since pantothenate was found in much lower abundance in *h2-1* (Dataset S1), we hypothesized that the lower level of TCA intermediates and the higher abundance of pyruvate could be explained by a shortage of CoA, which would thus reduce the flux from glycolysis to TCA cycle, and also potentially affects the conversion of α-ketoglutarate to succinyl-CoA;
3. Also associated with the GCS activity, the C1 cycle supports methionine biosynthesis. Since the abundance of methionine was lower in *h2-1*, we hypothesized that lower methionine levels would affect the pool of formylmethionine, which is a precursor for the translation of mitochondrially encoded proteins. Since protein complexes from the electron transport chain (ETC) require a coordinated expression and translation of proteins both nuclear and mitochondrially encoded, such scenario would imply an impairment of the ETC and an altered ATP production. In turn, this could affect the energy level of the cell, which would repress the progression of the cell cycle as already shown in (Baena-Gonzalez et al 2007; Mair et al 2015; Pedrotti et al 2018; Canal et al 2024).

In order to test all these hypotheses, we combined different approaches. First, we used several methods based on LC-MS to quantify key metabolites (Dataset S2). This revealed that T6P levels were actually found in higher abundance in *h2-1* than in WT root cells (Fig. 4d, SFig. 6), hence invalidating our first hypothesis. Secondly, CoA and acetyl-CoA levels were also found in higher abundance in roots of *h2-1* than of WT, while most of glycolytic intermediates remained unchanged (Fig. 4d, SFig. 6). Furthermore, the addition of CoA, acetyl-CoA, pantothenate or even ascorbate to *h2-1* seedlings grown on plates did not rescue their root growth (Fig. 4e; Dataset S3). Therefore, this ruled out our second hypothesis. Finally, while the pool of adenylates was unchanged (Fig. 4d), the addition of 10 or 50 μM of *f*-methionine slightly, but significantly, rescued the root growth of *h2-1*. However, immunoblots with antibodies raised against COX2 (cytochrome oxidase 2), a mitochondrial protein encoded by mitochondrial genome; IDH (isocitrate dehydrogenase), a mitochondrial protein encoded by the nuclear genome, and HIS3 (histone H3), a nuclear protein encoded by the nuclear genome, showed that the ratio between mitochondrially and nuclear encoded proteins did not differ between WT and mutant (Fig. 4f). Taken together, these results did not support our third hypothesis either, and prompted us to propose another theory.

**Figure 4.**
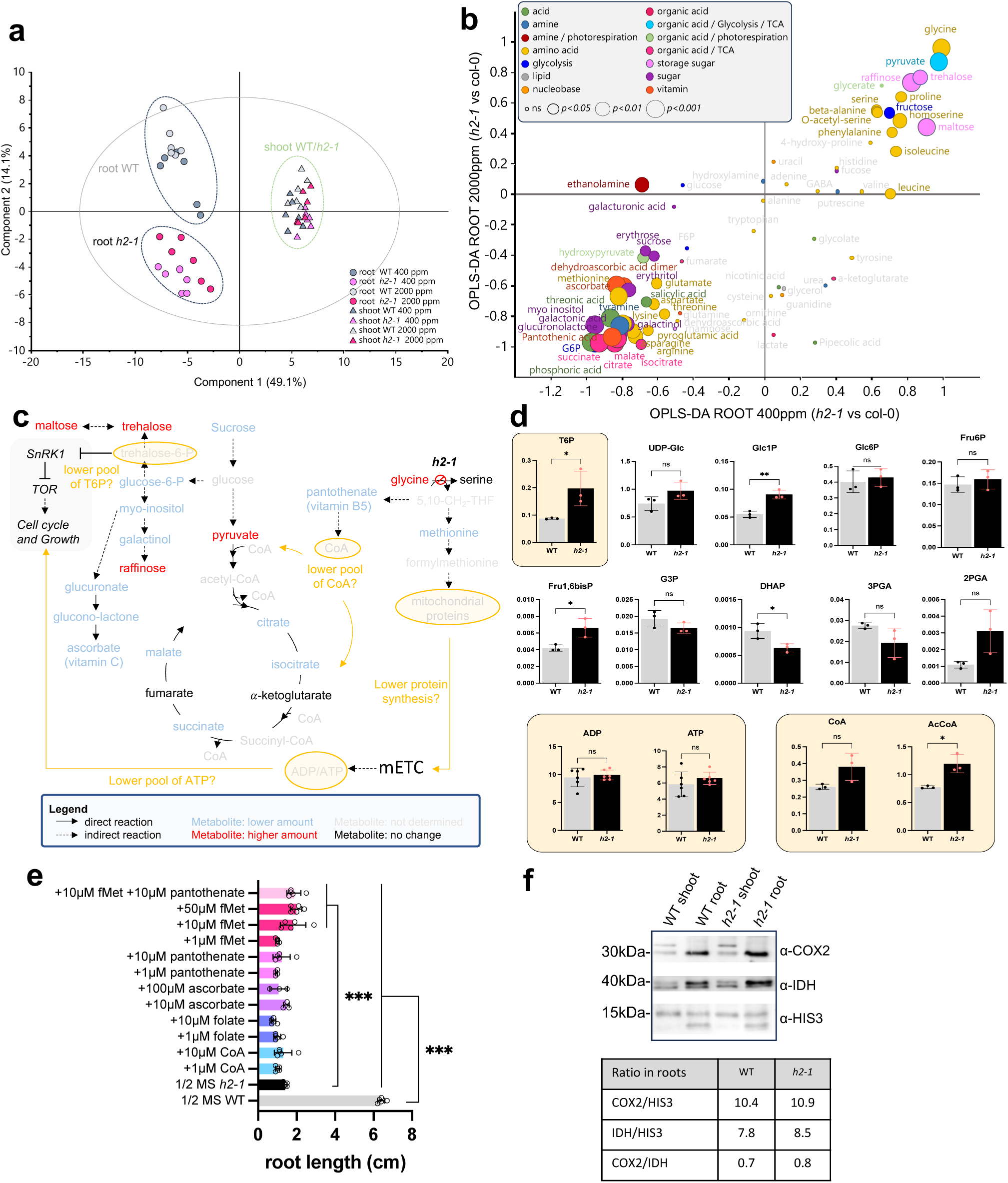
Metabolomics and bioassays. **(a)** PCA untargeted analysis WT vs *h2-1*, root/shoot samples from plans grown at either 400 ppm or 2000 ppm. (**b**) SUS-Plot root samples WT vs *h2- 1* at 400 and 2000 ppm. **(c**) Working model for altered primary metabolism in *h2-1* roots, and working hypotheses. For details see text. **(d)** Targeted analysis for key metabolites in WT and *h2-1* roots. (**e**) Root length of *h2-1* after 14 days on plates supplemented with compounds, WT root length in 1/2 MS is used as control. (**f**) Ratio between the abundance of mitochondrial proteins nuclear and mitochondrion encoded in WT and *h2-1* roots.

### Mitochondrial lipoylation in heterotrophic tissue is affected in *h2-1*

It has been shown that lipoylation in mitochondria of photosynthetic cells could occur independently of the H-protein, notably via a salvage system showing some similarities with that found in prokaryotes (Fig. 5a; Ewald et al 2007; Ewald et al 2014; Guan et al 2016; 2017). This salvage system requires a Lipoate-Protein Ligase (LPLA), an essential mitochondrial enzyme that would use octanoyl-nucleoside monophosphate, and possibly other donor substrates including lipoyl-AMP, for the octanoylation or lipoylation of mitochondrial PDH-E2, αKGDH-E2 and GCS-H proteins (Fig. 5b; Ewald et al 2014). Until now, it was assumed that such a lipoylation system would work for all cell types in plants. However, the accumulation of pyruvate and BCAA, the lower levels of TCA intermediates combined with all the other results described above prompted us to test whether the lipoylation system would work differently in heterotrophic tissue as compared to what has been reported so far for plants, and notably for photosynthetic cells. For this purpose, we performed immunoblot analyses using anti-lipoate antibodies on a total protein extract from shoots of WT and *h2-1* mutant plants (Fig. 5b). No differences in the lipoylation profile of target proteins could be observed for shoots. Intriguingly, a similar western blotting analysis with root samples gave inconsistent results, likely due to the very low amount of material as well as the contamination from photosynthetic cells from the hypocotyl. To circumvent this problem, we transformed WT and *h2-1* plants with an IMTACT construct (Isolation of Mitochondria TAgged in specific Cell Types; Boussardon et al 2020, Boussardon and Keech, 2022) allowing for the synthesis *in cellulo* of an extension eGFP-BLRP (Biotin Ligase Receptor Protein) located on the outer membrane protein OM64. This expression was driven by the promoter of *H2* (*pH2*) (Fig. 5c). Such a strategy allowed us to only trap mitochondria from WT and *h2-1* in which *H2* is spatio-temporally expressed. Immunoblots with anti-lipoate antibodies on crude extracts, i.e. samples enriched with mitochondria from the whole seedlings, and with purified mitochondria from root tissues only expressing *H2* revealed that *h2-1* had a severely compromised lipoylation profile as compared to WT samples. No lipoylation of E2-subunits of PDH, αKGDH, and perhaps of BCKDH, as well as of H2, logically, could be detected in purified mitochondria of *h2-1* roots (Fig. 5d). Interestingly, one band around 40kDa was still present in all IMTACT samples, while another band appeared at ca. 25kDa only in *pH2*-mitochondria from *h2-1* mutants.

**Figure 5.**
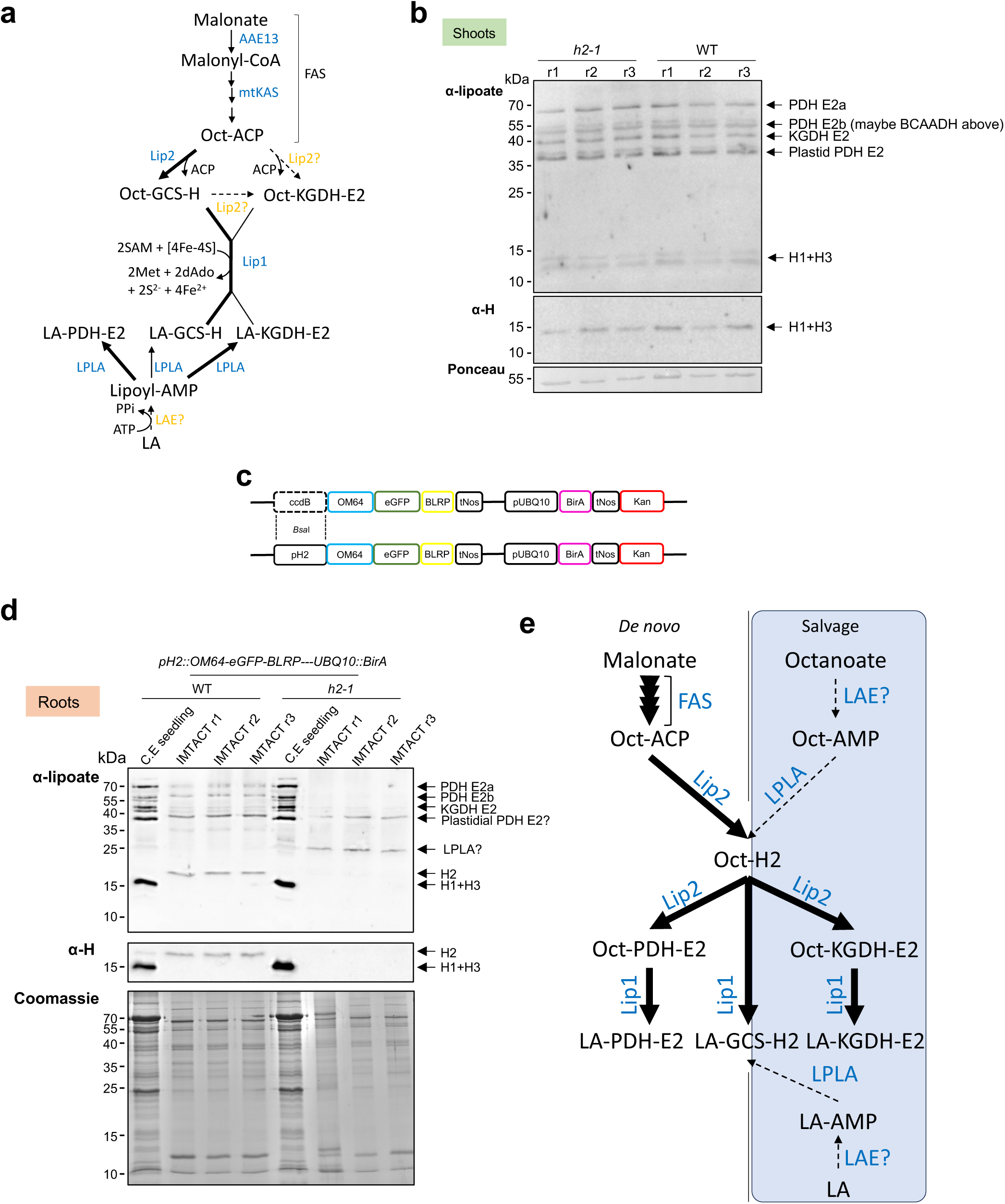
Current models for lipoylation in Arabidopsis and lipoylation state in root and shoots of WT and *h2-1*. (**a**) Current model for lipoylation in plants; for details see text. AAE13: malonyl-CoA synthetase, ACP: acyl carrier protein, AMP, adenosine monophosphate, FAS: fatty acid system, GCS-H2: H2 protein, KGDH: alpha-ketoglutarate dehydrogenase, LA: lipoic moiety, LAE: lipoate-activating enzyme, Lip1: lipoate synthase, Lip2: octanoyl transferase, LPLA: lipoate-protein ligase A, mtKAS: mitochondrial b-ketoacyl-acyl carrier protein synthase, Oct: octanoyl moiety, PDH: pyruvate dehydrogenase. **(b)** Lipoylation profile in shoot of Arabidopsis. WT and *h2-1* total extracts (10µg per well), r1-3 are biological replicates **(c)** IMTACT constructs used to transform WT and *h2-1* genotypes to isolate only mitochondria where H2 is expressed. **(d)** Lipoylation profile of mitochondria in roots (10µg for CE, and 3µg for IMTACT samples per well), CE: crude extract enriched with mitochondria from the whole seedling, IMTACT: only mitochondria where H2 is normally expressed. **(e)** Proposed model for mitochondrial lipoylation in root/heterotrophic cells. H2 acts as the main octanoyl donor for E2 mitochondrial proteins. A plain arrows indicates a demonstrated reaction whereas a dashed arrow indicates a still unclear step; thickness of the arrows represents the flux.

## Discussion

### Acquisition of specific spatiotemporal regulations for the metabolic use of H-proteins in land plants during evolution

Phylogeny analyses have pointed to an α-proteobacterial origin of the GCS subunits P, T and L in mitochondria of plants and algae (Kern et al 2011). Yet, the authors also mentioned that the phylogenetic relation of the GCS H-protein could not be finally resolved due to the rather short sequence, which made statistical save branching impossible. In this work, we show that mitochondrial H-proteins form two distinct phylogenetic clades in land plants: one that clusters with H1 and H3, and another that clusters with H2. This contrasts with what is observed in other eukaryotes (yeast, mammals), which possess a unique H representative. The absence of H2 in *h2-1* mutants results in a >99% impeded biomass production as compared to WT plants after 7 weeks in short day conditions. Since H2 is nearly solely present in heterotrophic tissues, it was no surprise to see that the growth phenotype was not rescued by non-photorespiratory conditions. Besides, the constitutive and ectopic expression of H1 or H3 fully restored the impeded growth phenotype of a *h2* mutant (Fig. 2), which implies that all three isoforms fulfill a similar metabolic function. Therefore, this suggests that land plants have acquired two specific spatiotemporal regulations for the metabolic use of H-proteins, and that the heterotrophic version of H-protein is essential for plant growth.

### Plant growth defaults of an h2-1 mutant could be partly attributed to a lower activation of TOR proteins in roots

Comparing WT and *h2-1* seedlings grown in light or in darkness (Fig. 3a-d) as well as using a reporter construct for the cell cycle progression, *pCycB1-eGFP* (Fig. 3e-f), allowed us to identify a defect in cell division in the root apical meristem of *h2-1*. In addition, a lower TOR kinase activity was detected in the roots of the *h2-1* mutant (Fig. 3g-h), confirming a down- regulation of the regulatory module controlling cell division and consequently growth. The TOR kinase complex is known to be repressed by another kinase, SnRK1 (Baena-González and Hanson, 2017), itself repressed by T6P (Zhang et al 2009; Nunes et al 2013). It was also shown that levels of T6P are linked to changes in carbon status; notably T6P closely follows endogenous levels of sucrose during diurnal cycles (Martins et al 2013). Intriguingly, we did not observe a lower abundance of T6P in the roots of *h2-1* as compared to WT roots (Fig. 4), which neither fits with the lower sucrose levels detected in *h2-1*, nor with a putative repression of TOR activity via the repression of SnRK1 by T6P. With that said, most of the data published for the T6P-SnRK1-TOR regulatory module were obtained from photosynthetic tissues. Furthermore, the actual repression of SnRK1 by T6P is still a matter of debate, and has been discussed expertly (Lunn et al 2014, Figueroa and Lunn 2016; Fichtner and Lunn 2021). For instance, it was recently shown that elevated T6P decreased *in vivo* SnRK1 activity in the light period, but not at the end of the night (Avidan et al, 2023), suggesting that SnRK1 activity is regulated by T6P in a context-dependent manner. So, although our data may in appearance conflict with the current model for photosynthetic tissues, they support the idea that the relationship between T6P and SnRK1 in developing tissues is complex and not yet fully resolved, particularly when it comes to tissue-specific regulations (Fichtner and Lunn 2021). Also worth mentioning, the metabolic shift observed in *h2-1* is another clear demonstration of the coupling between mitochondrial energy metabolism and TOR-dependent control of the cell cycle and growth (Siqueira et al 2018; O’Leary et al 2020; Canal et al 2024). However, while TCA intermediates were found in significantly lower abundance, the pools of adenylates (ATP and ADP) were not different between WT and *h2-1*, which raises the question about the actual signal(s) that can repress the TOR kinase activity at the cellular level.

### Autotrophic and heterotrophic tissues from land plants may have different mitochondrial lipoic acid metabolism

H-protein, which has no catalytic activity, plays a preponderant role in the GCS complex by serving as a mobile co-substrate for the three other enzymes. The mechanistic core of this complex relies on its lipoyl arm, which undergoes a catalytic cycle involving successive reductive methylamination, demethylamination and re-oxidation during glycine decarboxylation. In addition, other mitochondrial complexes, i.e. PDH, αKGDH and BCKDC, also require lipoylation for their catalytic activity (Reed 1974; Taylor et al 2004), where lipoic acid is covalently bound to specific Lys residues of the dihydrolipoyl transacetylase (E2) subunits of the complexes. In turn, E2 couples the decarboxylase (E1) and dihydrolipoyl dehydrogenase (E3) subunits, providing a means of acyl group movement to CoA and a mechanism of electron transfer to synthesize NADH. Here, it is worth mentioning that besides E2, E3 subunits can also be lipoylated (Taylor et al 2004).

Eukaryotes and prokaryotes have slightly diverging lipoic acid metabolism. Prokaryotes possess two clearly identified pathways for protein lipoylation: a *de novo* lipoic acid synthesis and a salvage system *per se*. Yet, even among prokaryotes, small differences have been reported. For instance, in *E. coli*, only two enzymes are required (SFig. 7): LipB, an octanoyl transferase, transfers an octanoyl moiety from octanoyl-ACP to each of the cognate protein substrates; then LipA, a lipoyl synthase, uses S-adenosyl-L-methionine and inserts two sulfur atoms to produce dihydrolipoyl moieties. A more complex pathway is found in *Bacillus subtilis* and *Staphyloccus aureus*, for which lipoyl moieties are first assembled on the GcvH protein (i.e. H-protein ortholog), and then transferred to the lipoyl domains of the α-ketoacid dehydrogenases. This pathway requires a lipoyl amidotransferase called LipL and a distinct octanoyl transferase called LipM (SFig. 7; Cao et al 2018). Interestingly, the current model for the *de novo* pathway of the synthesis and transfer of lipoic acid in humans, yeast and in Arabidopsis shows similarities to the one described for *Bacillus subtilis* and *Staphyloccus aureus*, even though a few divergences have been reported (SFig. 7; Mayr et al 2014; Cao et al 2018; Solmonson and DeBerardinis 2018). For instance, *S. cerevisiae* has both pathways in their mitochondria, while to date, no LPLA has been reported in *H. sapiens*, which thus argues against the presence of a salvage pathway (Solmonson and DeBerardinis 2018; Cao et al 2018). In Arabidopsis, both a *de novo* and a salvage system are present (Fig 5a; Ewald et al, 2007; Ewald et al 2014; Guan et al 2016; 2017). Yet, several significant divergences with other organisms are noted, which seems to make the plant model unique. Firstly, no lipoyl amidotransferase (Lip3; SFig. 7) has been identified in Arabidopsis. Secondly, it was reported that the deletion of mtKAS (mitochondrial βketoacyl-[acyl carrier protein] synthase), an enzyme of the mitochondrial fatty acid system (FAS) that provides the octanoyl-ACP for the octanoylation of H-protein, did affect the lipoylation status of mitochondrial proteins in shoots of Arabidopsis (Ewald et al, 2007; Guan et al 2017); however, the *mtkas* mutant did not show any apparent effect on mitochondrial lipoylation of PDH-E2 and αKGDH-E2 in roots (Fig. 5a; Ewald et al, 2007). Similarly, Guan et al (2016) showed that a knockout of AAE13, a Malonyl-CoA synthetase part of FAS, also nearly abolished lipoylation of H-protein in Arabidopsis shoots, but had a much less severe effect on PDH and αKGDH. Thirdly, even though an ortholog of *E. coli* Lipoate-Protein Ligase (AtLPLA) was reported in Arabidopsis, and supports the presence of a salvage pathway for lipoylation in plants (Ewald et al 2014), the substrate for AtLPLA remains unclear (Fig. 5a). In addition, the authors showed that in Arabidopsis roots, lipoylation of H-protein was not affected in *RNAi-LPLA* lines whereas it was strongly affected in both PDH-E2 and αKGDH-E2 (Ewald et al 2014). Considered together with the results of this study, this suggests that lipoylation in mitochondria from autotrophic and heterotrophic tissues, i.e. shoot and roots cells here, may follow different mechanisms. Therefore, we propose a novel model where H2 is the main relay for lipoylation in mitochondria from root cells (Fig 5e). This model diverges from the current understanding of lipoylation in plant cells, and besides positioning H2 as a key donor in a similar manner as many other organisms, it also raises new questions. For example, why AtLPLA does not, at least partially, rescue the mitochondrial lipoylation in *h2* mutant line? At this stage, we can suggest two possibilities: 1) AtLPLA in roots takes only lipoate from H-protein, and 2) the pool of lipoate available might be limiting since there is no H-protein for its biosynthesis, so in turn there might be too little lipoate to rescue. Yet, additional research effort is needed to decipher the different pathways for lipoylation in plant mitochondria, and determine the extent to which lipoylation differs between autotrophic and heterotrophic tissues.

## MATERIALS AND METHODS

### Plant material and growth conditions

Arabidopsis T-DNA insertion lines were obtained from the Arabidopsis Biological Resource Center: *h2-1* (SALK_120199; Col-0); *h2-2* (SAIL_1152_G07; Col-0). Homozygous plants were selected by genotyping PCR and primers used are reported in the Table S1. Seeds were sown directly on mixture soil:vermiculite 3:1 or were surface sterilized using successive ethanol solution at 70% and 100% before spreading on ½ Murashige and Skoog (MS)-agar 0.7% (w/v) medium supplemented with 0.1% sucrose.

After vernalization for 48 h at 4°C in darkness, plants were grown under short-day photoperiod (SD: light 8h at 22°C, dark 16h at 17°C) at a photosynthetically active radiation of 180 μmol m^-2^ s^-1^ and a relative air humidity of 65% or under long-day photoperiod (LD: light 16h at 22°C, dark 8h at 17°C) at a photosynthetically active radiation of 150 μmol m^-2^ s^-1^ and a relative air humidity of 65%.

For hydroponic growth system, the protocol provided by Conn et al 2013 was followed. Basically, sterilized Arabidopsis seeds were sown on germination medium-agar 0.7% contained in black microcentrifuge lids. The culture system was covered with plastic clingfilm during 1 week to enhance humidity. After 2 days in the dark at 4°C, stratified seeds were transferred under long-day photoperiod as mentioned above. After 7 days, germination medium was replaced by a modified ¼ Hoagland’s solution referred to basal nutrient solution. Then, the basal nutrient solution was weekly replaced until analysis.

### Evolutionary and sequence analysis

Protein sequence were retrieved from The Arabidopsis Information Resource (TAIR). Alignment was obtained by using ClustalW (BioEdit) program. Phylogenetic tree was constructed with MEGA7 software and using the Neighbor-Joining method.

### Gene expression analysis

Samples were frozen in liquid N^2^ in 2 mL tubes containing two steel beads (2.5 mm diameter). Tissues were disrupted for 1 min at 30 s^-1^ in a Retsch mixer mill MM301 homogenizer (Retsch, Haan, Germany). Total RNA was extracted from 2-week-old seedlings using the E.Z.N.A.® Plant RNA kit (Omega bio-tek). Extracted RNA were then treated by DNAse treatment (ThermoFischer, DNAseI). The integrity of RNA was systematically verified on agarose gel.

Total RNA was quantified with a Nanodrop (ThermoFischer; Nanodrop^TM^ 2000). Reverse transcription was achieved from 1 µg of RNA using SuperScript™ III Reverse Transcriptase (ThermoFischer). RT-PCR reactions were performed in a total volume of 20 µL. Amplification reactions were performed in a Bio-Rad CFX-96 real-time PCR system. The house-keeping gene *ACTIN-2* (*At3g18780*) was used as reference gene.

### Protein extraction and immunoblot analyses

Total protein extracts were prepared using a protein extraction buffer (100 mM Tris-HCl pH 7.5, 50 mM EDTA, 250 mM NaCl, 0.05% SDS). Protein quantification was done with a Bradford protein assay (Bio-Rad). Soluble proteins were separated on SDS-PAGE 12% and transferred to nitrocellulose membrane. Membranes were blocked with 5% milk in PBS-T for 1 h, followed by an overnight incubation at 4°C with specific polyclonal primary antibodies raised against GCS-H proteins (1/1000, Agrisera, AS05 074), the nuclear histone 3 (HIS3, 1/5000, Agrisera, AS10 710), mitochondrial isocitrate dehydrogenase protein (IDH, 1/1000, Agrisera, AS06 203A), plant cytochrome oxidase subunit II (COXII, Agrisera, AS04 053A), Lipoic Acid (LA, 1/1000, Merck), RPS6 (1/1000, Cell Signaling Technology, 2217S), and RPS6P (1/2000; Dobrenel et al 2016), diluted in 2 % milk in PBS-T. After 2 h incubation at room temperature with anti-rabbit IgG conjugated to horseradish peroxidase (1/2000 or 1/10000 in 2 % milk in PBS-T, Agrisera, AS09 602) or anti-mouse in the case of RPS6 (1/2000 in 2 % milk in PBS-T, Agrisera, AS11 1772). Signals were detected by chemiluminescence (Agrisera ECL kit bright; AS16 488 ECL-N-100) using Azure c600 Western Blot Imaging system (Azure biosystems). Exposure time was 1-5 minutes. Ponceau S staining (0.2% ponceau S, 3% trichloroacetic acid) of total proteins was systematically performed after revelation.

#### RPS6 phosphorylation status

To study RPS6 phosphorylation, 10 µg (roots) and 20 µg (shoots) of proteins per well were loaded. Membranes were blocked with 5% milk in PBS-T for 1 h followed by an overnight incubation at 4°C with specific polyclonal primary antibodies anti- RPS6 (1/1000, Cell Signaling Technology, 2217S), anti-RPS6P (1/2000; Dobrenel et al 2016), diluted in 2 % milk in PBS-T. After 2 h incubation at room temperature with a goat anti-rabbit (for RPS6P) or a goat anti-mouse (for RPS6) secondary antibodies conjugated to horseradish peroxidase (1/2000 in 2 % milk in PBS-T, Agrisera, AS09 602 and AS11 1772). Visualization and exposure time were as described above. Gels replicates are shown in SFig. 5.

### Multivariate analysis

“Multivariate analysis (MVA) was performed using the SIMCA software (v.17.0.2.34594) [www.sartorius.com/umetrics]. Principal component analysis (PCA) was used for unsupervised modelling and orthogonal partial least squares discriminant analysis (OPLS-DA) was used for supervised classification (Bylesjö et al 2006). The shared and unique structure (SUS) plot was used for metabolite contribution comparisons between models (Wiklund et al 2008).

### IMTACT constructs and isolation of mitochondria

One-shot plasmid *ccdb::OM64-eGFP-BLRP UBQ10::BirA* were obtained from Boussardon et al. (2020). The *H2* promoter (1420bp) was amplified by PCR (adding *BsaI* sites in 5’ and 3’) to create LI PCR products. The *ccdB* cassette was replaced by *pH2* using a LI/LIII ligation/cut reaction (100 ng LIII vector, 5 µl of purified PCR product, 1.5 µl 10X Buffer G, 1.5 µl 10mM ATP, 0.75 µl BsaI 20 U/µl, 0.75 µl T4 DNA ligase 5 U/µl, H^2^O to 15 µl).

Plasmids were used for *Agrobacterium tumefaciens* (GV3101::pMP90, pSOUP) transformation of Col-0 using the floral dip method (Clough and Bent, 1998). *In vitro* selections of T1 plantlets were made on ½ MS + 0.1% sucrose media containing Basta (10 μg/ml).

A detailed IMTACT procedure can be found in Boussardon and Keech (2022). Briefly, a crude extract was obtained from leaves or roots and incubated for 5 minutes with 30 µl of magnetic beads (ThermoFischer, Dynabead MyOne Streptavidin T1, 65601). After washing, mitochondria couple to beads were resuspended with 50 µl of wash buffer. Protein quantification was done with a Bradford protein assay (Bio-Rad).

### GUS activity

β-glucuronidase (GUS) fusion plasmids were prepared by replacing the CaMV 35S promoter in vector pCAMBIA 1305.1 with about 1 kb promotor fragments from the respective Arabidopsis H-protein genes including their 5’-untranslated regions. Seedlings and organs of transformed Arabidopsis were stained for GUS expression essentially as described in Vitha et al (1995).

### Subcellular localization

#### Cell cycle

Fluorescence of the eGFP protein under the control of pcycB1 was observed on a confocal microscope (Zeiss, LSM780) and a Zeiss Zen software. Eight-day-old Col-0 and H2 plantlets grown on a ½ MS (0.8% agarose) media were collected and signals were detected according to the following excitation/emission wavelengths: eGFP (488 nm/495-535 nm; gain: 750). Pictures were analysed using the ImageJ software (https://imagej.nih.gov/ij/).

#### Root mitochondria

Eight-day-old Col-0 and H2 grown on a ½ MS (0.8% agarose) media were vacuum infiltrated with 100 nM of Mitotracker Orange CMTMRos (ThermoFischer, M7510), diluted in ½ MS, for 5 min and incubated 15 min in darkness. Signals were detected according to the following excitation/emission wavelengths: Mitotracker (561 nm/575-630 nm; gain: 500). Pictures were analysed using the ImageJ software (https://imagej.nih.gov/ij/).

### Plasmid editing

As an *h2-1* mutant is kanamycin resistant, the *pSL36 pcycB1::eGFP* vector was edited to introduce a gene conferring a resistance to hygromycin. To this aim, the hygromycin resistance gene of the *pUB-DEST* destination vector (UBQ10 promoter, no tag), flanked by *Sac*I in 5’ and *Xba*I restriction sites in 3’, was amplified. Briefly, 3 parts were amplified using (i) SacI_pSL36- F and pSL36_Hygro-R, (ii) Hygro_pSL36-F and Hygro_pSL36-R, (iii) pSL36_Hygro-F and XbaI_pSL36-R. The 3 amplicons were combined with SacI_pSL36-F and XbaI_pSL36-R. SacI and XbaI restriction assay was performed in pSL36 pcycB1::eGFP----Kan plasmid and the hygromycin combined PCR product. Hence, the hygromycin amplicon (3899 bp) was used to replace the kanamycin containing region of pSL36 (3671 bp). The hygromycin gene is under the control of the NOS promoter, present in pSL36. Sequence was verified by Sanger sequencing.

### Bioassays

*h2-1* and WT Col-0 seeds were sterilized and stratified for 3 days and sown on ½ MS supplemented with formyl-methionine (1, 10 or 50 µM), coenzyme A (1 or 10 µM), folate (1 or 10 µM), vitamin B5 (1 or 10 µM), ascorbate (1 or 10 mM), formyl-methionine and vitamin B5 (10 µM each or 50 µM and 10 µM). Root growth was monitored every two days for 14 days in a long-day growth chamber.

### Metabolomic analysis

*GC-MS metabolites profiling (untargeted analysis)* – Extraction and analysis by gas chromatography–mass spectrometry (GC-MS) was performed as described in Lisec et al. (2006). Briefly, plant material was extracted in 700 μl of methanol at 70 °C for 15 min and 375 μl of chloroform followed by 750 μl of water was added. The polar fraction was dried under vacuum, and the residue was derivatized for 120 min at 37 °C (in 40 μl of 20 mg ml−1 methoxyamine hydrochloride (Sigma-Aldrich, cat. no. 593-56-6) in pyridine followed by a 30 min treatment at 37 °C with 70 μl of N-methyl-N-(trimethylsilyl)trifluoroacetamide (MSTFA reagent; Macherey-Nagel, cat. no. 24589-78-4). An autosampler Gerstel Multi-Purpose system (Gerstel GmbH & Co.KG, Mülheim an der Ruhr, Germany) was used to inject 1µl of the samples in splitless mode to a chromatograph coupled to a time-of-flight mass spectrometer system (Leco Pegasus HT TOF-MS; Leco Corp., St Joseph, MI, USA). Helium was used as carrier gas at a constant flow rate of 2 ml s−1 and GC was performed on a 30 m DB-35 column (capillary column, 30 m length, 0.32 mm inner diameter, 0.25 μm film thickness, PN: G42, Agilent). The injection temperature was 230 °C and the transfer line and ion source were set to 250 °C. The initial temperature of the oven (85 °C) increased at a rate of 15 °C min−1 up to a final temperature of 360 °C. After a solvent delay of 180 s, mass spectra were recorded at 20 scans s−1 with m/z 70–600 scanning range. Chromatograms and mass spectra were evaluated using Chroma TOF 4.5 (Leco) and Xcalibur 4.0 (Thermo) software. Metabolites were annotated based on a retention time compared with the reference data as described in (Alseekh et al 2021).

#### ATP/ADP and Acetyl-CoA/CoA

Approximately 15 mg of plant tissue were extracted as follows: 360 µL of 80% methanol containing 0.1 M formic acid were added to the samples on ice, immediately after addition of extraction solvent the samples were vortexed for 5 seconds and 20 µL of 9% ammonium bicarbonate (*w*/*v* in water) were added; the samples were vortexed again for 5 seconds and each sample was shaken with a tungsten carbide bead at 30 kHz for 3 minutes, then the samples were centrifuged at 14000 g for 10 minutes. The supernatant was transferred to LC vials and dried using a centrifugal concentrator (Thermo Fisher Scientific, Waltham, USA), and finally reconstituted in 50 µL of 50% methanol. Subsequently, the samples were analyzed using liquid chromatography-tandem mass spectrometry (LC-MS/MS) system consisting of an Agilent 1290 UPLC connected to an Agilent 6490 triple quadrupole (Agilent, CA, USA), metabolites were detected in multiple-reaction monitoring (MRM) mode.

The separation of ATP/ADP and Acetyl-CoA/CoA were achieved by injecting 3 µL of the extracts to a HILIC column (iHILIC-(P) Classic, PEEK, 50 × 2.1 mm, 5 µm, HILICON, Umeå, Sweden), the column and autosampler were maintained at 40 °C and 4 °C, respectively. The mobile phase composed of (A) 10 mM ammonium acetate with 5 µM medronic acid in water of pH 6.8 and (B) 10 mM ammonium acetate in 90% acetonitrile. The mobile phase was delivered at a flow rate of 0.35 mL/min at the following gradient elution: 0.0 min (85% B), 5 min (60% B), 7 min (30% B), 8 min (30% B), 9 min (85% B), 15 min (85%). Analytes were ionized in electrospray source operated in the positive mode. The source and gas parameters were set as follows: ion spray voltage 4.0 kV, gas temperature 200 °C, drying gas flow 11 L/min, nebulizer pressure 30 psi, sheath gas temperature 375 °C, sheath gas flow 12 L/min, fragmentor 380 V. The instrument was operated in dynamic multiple reaction monitoring mode (MRM), and the MRM transitions of metabolites of interest were optimized by using Agilent MassHunter Optimizer.

#### Sugar phosphates

Approximately 10 mg of each sample was extracted in 1 mL cold extraction mixture consisting of chloroform:MeOH:H2O (1:3:1). To each sample, 1 µg of 2- deoxy-d-glucose 6-phosphate was added as internal standard. Samples were treated in a mixer mill set to a frequency 30 Hz for 3 min, with 2 tungsten carbide beads added to each tube. Obtained extracts were centrifuged at 14000 g for 10 min. 250 µL of the supernatant were transferred into an LC vial followed by evaporation with a SpeedVac. For derivatization, dried samples were dissolved in 20 µL of methoxylamine and incubated on a heat block at 60 °C for 30 min. After incubation at room temperature overnight, 6 µL of 1-methylimidazole and 12 µL of propionic acid anhydride were added and heated at 37 °C for 30 min. The reaction mixture was then evaporated to dryness by N2 gas. Prior to LC–MS analysis, derivatized metabolites were dissolved in 100 µL of aqueous 0.1% formic acid. Quantitative analysis was done by combined ultrahigh-performance liquid chromatography-electrospray ionization-triple quadrupole-tandem mass spectrometry (UHPLC-ESI-QqQ-MS/MS) in dynamic multiplereaction-monitoring (MRM) mode. An Agilent 6495 UHPLC chromatograph equipped with a Waters Acquity BEH 1.7 µm, 2.1×100 mm column (Waters Corporation, Milford, USA) coupled to a QqQ-MS/MS (Agilent Technologies, Atlanta, GA, USA) was used. The washing solution for the auto sampler syringe and injection needle was 90% MeOH with 1% HCOOH. The mobile phase consisted of A, 2% HCOOH and B, MeOH with 2% HCOOH. The gradient was 0% B for 1 min followed by linear gradients from 0.1 to 30% B from 1 to 3 min then 30–40% B from 3 to 6 min, hold at 40% B from 6 to 10 min, followed by 40–70% B from 10 to 12.5 min, hold at 70% B from 12.5 to 15 min, and thereafter 70–99% B from 15 to 17.5 min. B was held at 99% for 0.5 min, and thereafter the column was re-equilibrated to 0% B. The flow rate was 0.65 mL.min^−1^ during equilibration and 0.5 mL.min^−1^ during the chromatographic runs. The column was heated to 40 °C, and injection volumes were 1 μL. The mass spectrometer was operated in negative ESI mode with gas temperature 230 °C; gas flow 12 L.min^−1^; nebulizer pressure 20 psi; sheath gas temperature 400 °C; sheath gas flow 12 L.min^−1^ ; capillary voltage 4000 V (neg); nozzle voltage 500 V; iFunnel high pressure RF 150V; iFunnel low pressure RF 60V. The fragmentor voltage 380V and cell acceleration voltage 5V. Data were processed using MassHunter Qualitative Analysis and Quantitative Analysis (QqQ; Agilent Technologies, Atlanta, GA, USA) and Excel (Microsoft, Redmond, Washington, USA) software.

### Accession Numbers

Accession numbers of genes studied in this work are as followed *H1: At2g35370; H2: At2g35120; H3: At1g32470*.

## Supporting information

Table S1_Primers

## Acknowledgements

We thank Dr Fazeelat Karamat for helping and sharing materials for hydroponic systems; the Swedish Metabolomic Center for their analyses; and Dr Thomas Dobrenel for his help with the RPS6P assays and critical discussion regarding the T6P-dependent regulations of TOR; Sylvie Figuet for technical assistance with plant growth.

## References

Alonso JM, Stepanova AN, Leisse TJ, Kim CJ, Chen HM, Shinn P, Stevenson DK, Zimmerman J, Barajas P, Cheuk R, Gadrinab C, Heller C, Jeske A, Koesema E, Meyers CC, Parker H, Prednis L, Ansari Y, Choy N, Deen H, Geralt M, Hazari N, Hom E, Karnes M, Mulholland C, Ndubaku R, Schmidt I, Guzman P, Aguilar-Henonin L, Schmid M, Weigel D, Carter DE, Marchand T, Risseeuw E, Brogden D, Zeko A, Crosby WL, Berry CC, Ecker JR (2003) Genome-wide Insertional mutagenesis of Arabidopsis thaliana. Science 301: 653–657

Alseekh S, Aharoni A, Brotman Y, Contrepois K, D’Auria J, Ewald J, J CE, Fraser PD, Giavalisco P, Hall RD, Heinemann M, Link H, Luo J, Neumann S, Nielsen J, Perez de Souza L, Saito K, Sauer U, Schroeder FC, Schuster S, Siuzdak G, Skirycz A, Sumner LW, Snyder MP, Tang H, Tohge T, Wang Y, Wen W, Wu S, Xu G, Zamboni N, Fernie AR (2021) Mass spectrometry-based metabolomics: a guide for annotation, quantification and best reporting practices. Nat Methods 18: 747–756

Avidan O, Moraes TA, Mengin V, Feil R, Rolland F, Stitt M, Lunn JE (2023) In vivo protein kinase activity of SnRK1 fluctuates in Arabidopsis roseTes during light-dark cycles. Plant Physiol 192: 387–408

Baena-González E, Hanson J (2017) Shaping plant development through the SnRK1-TOR metabolic regulators. Curr Opin Plant Biol 35: 152–157

Baena-Gonzalez E, Rolland F, Thevelein JM, Sheen J (2007) A central integrator of transcription networks in plant stress and energy signalling. Nature 448: 938–942

Bauwe H, Hagemann M, Fernie AR (2010) Photorespiration: players, partners and origin. Trends in Plant Science 15: 330-336

Blackwell RD, Murray AJ, Lea PJ (1990) Photorespiratory mutants of the mitochondrial conversion of glycine to serine. Plant Physiol 94: 1316–1322

Boussardon C, Keech O (2022) Cell Type-Specific Isolation of Mitochondria in Arabidopsis. Methods Mol Biol 2363: 13–23

Boussardon C, Przybyla-Toscano J, Carrie C, Keech O (2020) Tissue-specific isolation of Arabidopsis/plant mitochondria - IMTACT (isolation of mitochondria tagged in specific cell types). Plant J 103: 459–473

Bylesjö M, Rantalainen M, Cloarec O, Nicholson JK, Holmes E, Trygg J (2006) OPLS discriminant analysis: combining the strengths of PLS-DA and SIMCA classification. Journal of Chemometrics 20: 341–351

Canal MV, Mansilla N, Gras DE, Ibarra A, Figueroa CM, Gonzalez DH, Welchen E (2024) Cytochrome c levels affect the TOR pathway to regulate growth and metabolism under energy-deficient conditions. New Phytologist 241: 2039–2058

Cao X, Zhu L, Song X, Hu Z, Cronan JE (2018) Protein moonlighting elucidates the essential human pathway catalyzing lipoic acid assembly on its cognate enzymes. Proc Natl Acad Sci U S A 115: E7063–e7072

Carrie C, Venne AS, Zahedi RP, Soll J (2015) Identification of cleavage sites and substrate proteins for two mitochondrial intermediate peptidases in Arabidopsis thaliana. J Exp Bot 66: 2691–2708

Clough SJ, Bent AF (1998) Floral dip: a simplified method for Agrobacterium-mediated transformation of Arabidopsis thaliana. Plant Journal 16: 735–743

Cohen-Addad C, Pares S, Sieker L, Neuburger M, Douce R (1995) The lipoamide arm in the glycine decarboxylase complex is not freely swinging. Nature Structural Biology 2: 63–68

Conn SJ, Hocking B, Dayod M, Xu B, Athman A, Henderson S, Aukett L, Conn V, Shearer MK, Fuentes S, Tyerman SD, Gilliham M (2013) Protocol: optimising hydroponic growth systems for nutritional and physiological analysis of Arabidopsis thaliana and other plants. Plant Methods 9: 4

Dobrenel T, Mancera-Marcnez E, Forzani C, Azzopardi M, Davanture M, Moreau M, Schepetilnikov M, Chicher J, Langella O, Zivy M, Robaglia C, Ryabova LA, Hanson J, Meyer C (2016) The Arabidopsis TOR Kinase Specifically Regulates the Expression of Nuclear Genes Coding for Plastidic Ribosomal Proteins and the Phosphorylation of the Cytosolic Ribosomal Protein S6. Frontiers in Plant Science 7

Douce R, Bourguignon J, Neuburger M, Rebeille F (2001) The glycine decarboxylase system: a fascinating complex. Trends in Plant Science 6: 167–176

Ducker GS, Rabinowitz JD (2017) One-Carbon Metabolism in Health and Disease. Cell Metab 25: 27–42

Engel N, van den Daele K, Kolukisaoglu U, Morgenthal K, Weckwerth W, Parnik T, Keerberg O, Bauwe H (2007) Deletion of Glycine Decarboxylase in Arabidopsis Is Lethal under Nonphotorespiratory Conditions. Plant Physiol. 144: 1328–1335

Ewald R, Hoffmann C, Florian A, Neuhaus E, Fernie AR, Bauwe H (2014) Lipoate-Protein Ligase and Octanoyltransferase Are Essential for Protein Lipoylation in Mitochondria of Arabidopsis. Plant Physiology 165: 978–990

Ewald R, Kolukisaoglu Un, Bauwe U, Mikkat S, Bauwe H (2007) Mitochondrial Protein Lipoylation Does Not Exclusively Depend on the mtKAS Pathway of de Novo FaTy Acid Synthesis in Arabidopsis. Plant Physiology 145: 41–48

Fichtner F, Lunn JE (2021) The Role of Trehalose 6-Phosphate (Tre6P) in Plant Metabolism and Development. Annu Rev Plant Biol 72: 737–760

Figueroa CM, Lunn JE (2016) A Tale of Two Sugars: Trehalose 6-Phosphate and Sucrose. Plant Physiology 172: 7–27

Fuchs P, Rugen N, Carrie C, Elsässer M, Finkemeier I, Giese J, Hildebrandt TM, Kühn K, Maurino VG, Ruberti C, Schallenberg-Rüdinger M, Steinbeck J, Braun H-P, Eubel H, Meyer EH, Müller-Schüssele SJ, Schwarzländer M (2020) Single organelle function and organization as estimated from Arabidopsis mitochondrial proteomics. The Plant Journal 101: 420–441

Fujiwara K, Okamura K, Motokawa Y (1979) Hydrogen carrier protein from chicken liver: purification, characterization, and role of its prosthetic group, lipolic acid, in the glycine cleavage reaction. Arch Biochem Biophys 197: 454–462

Guan X, Nikolau BJ (2016) AAE13 encodes a dual-localized malonyl-CoA synthetase that is crucial for mitochondrial faTy acid biosynthesis. The Plant Journal 85: 581–593

Guan X, Okazaki Y, Lithio A, Li L, Zhao X, Jin H, Nettleton D, Saito K, Nikolau BJ (2017) Discovery and Characterization of the 3-Hydroxyacyl-ACP Dehydratase Component of the Plant Mitochondrial FaTy Acid Synthase System. Plant Physiol 173: 2010–2028

Hanson AD, Roje S (2001) ONE-CARBON METABOLISM IN HIGHER PLANTS. Annual Review of Plant Physiology and Plant Molecular Biology 52: 119

Kern R, Bauwe H, Hagemann M (2011) Evolution of enzymes involved in the photorespiratory 2- phosphoglycolate cycle from cyanobacteria via algae toward plants. Photosynthesis Research 109: 103–114

Kikuchi G, Hiraga K (1982) The mitochondrial glycine cleavage system. Molecular and Cellular Biochemistry 45: 137–149

Kobayashi K, Suzuki T, Iwata E, Nakamichi N, Suzuki T, Chen P, Ohtani M, Ishida T, Hosoya H, Müller S, Leviczky T, Pettkó-Szandtner A, Darula Z, Iwamoto A, Nomoto M, Tada Y, Higashiyama T, Demura T, Doonan JH, Hauser MT, Sugimoto K, Umeda M, Magyar Z, Bögre L, Ito M (2015) Transcriptional repression by MYB3R proteins regulates plant organ growth. Embo j 34: 1992–2007

Lee CP, Eubel H, O’Toole N, Millar AH (2008) Heterogeneity of the Mitochondrial Proteome for Photosynthetic and Non-photosynthetic *Arabidopsis* Metabolism. Molecular & Cellular Proteomics 7: 1297–1316

Lisec J, Schauer N, Kopka J, Willmitzer L, Fernie AR (2006) Gas chromatography mass spectrometry-based metabolite profiling in plants. Nat Protoc 1: 387–396

Lunn JE, Delorge I, Figueroa CM, Van Dijck P, Stitt M (2014) Trehalose metabolism in plants. The Plant Journal 79: 544–567

Mair A, Pedroi L, Wurzinger B, Anrather D, Simeunovic A, Weiste C, Valerio C, Dietrich K, Kirchler T, Nägele T, Vicente Carbajosa J, Hanson J, Baena-González E, Chaban C, Weckwerth W, Dröge-Laser W, Teige M (2015) SnRK1-triggered switch of bZIP63 dimerization mediates the low-energy response in plants. eLife 4: e05828

Martins MC, Hejazi M, Fettke J, Steup M, Feil R, Krause U, Arrivault S, Vosloh D, Figueroa CM, Ivakov A, Yadav UP, Piques M, Metzner D, Stitt M, Lunn JE (2013) Feedback inhibition of starch degradation in Arabidopsis leaves mediated by trehalose 6-phosphate. Plant Physiol 163: 1142–1163

Mayr JA, Feichtinger RG, Tort F, Ribes A, Sperl W (2014) Lipoic acid biosynthesis defects. J Inherit Metab Dis 37: 553–563

Mukha D, Fokra M, Feldman A, Sarvin B, Sarvin N, Nevo-Dinur K, Besser E, Hallo E, Aizenshtein E, Schug ZT, Shlomi T (2022) Glycine decarboxylase maintains mitochondrial protein lipoylation to support tumor growth. Cell Metabolism 34: 775–782.e779

Nunes C, Primavesi LF, Patel MK, Martinez-Barajas E, Powers SJ, Sagar R, Fevereiro PS, Davis BG, Paul MJ (2013) Inhibition of SnRK1 by metabolites: tissue-dependent effects and cooperative inhibition by glucose 1-phosphate in combination with trehalose 6-phosphate. Plant Physiol Biochem 63: 89–98

O’Leary BM, Oh GGK, Lee CP, Millar AH (2020) Metabolite Regulatory Interactions Control Plant Respiratory Metabolism via Target of Rapamycin (TOR) Kinase Activation. Plant Cell 32: 666–682

Oliver D (1994) The Glycine Decarboxylase Complex from Plant Mitochondria. Annual Review of Plant Physiology and Plant Molecular Biology 45: 323–337

Oliver DJ, Neuburger M, Bourguignon J, Douce R (1990) Glycine metabolism by plant mitochondria. Physiologia Plantarum 80: 487–491

Pares S, Cohen-Addad C, Sieker L, Neuburger M, Douce R (1994) X-ray structure determination at 2.6-A resolution of a lipoate-containing protein: the H-protein of the glycine decarboxylase complex from pea leaves. Proc Natl Acad Sci U S A 91: 4850–4853

Pedroi L, Weiste C, Nagele T, Wolf E, Lorenzin F, Dietrich K, Mair A, Weckwerth W, Teige M, Baena-Gonzalez E, Droge-Laser W (2018) Snf1-RELATED KINASE1-Controlled C/S1-bZIP Signaling Activates Alternative Mitochondrial Metabolic Pathways to Ensure Plant Survival in Extended Darkness. Plant Cell 30: 495–509

Reed LJ (1974) Multienzyme complexes. Accounts of Chemical Research 7: 40–46

Ren M, Venglat P, Qiu S, Feng L, Cao Y, Wang E, Xiang D, Wang J, Alexander D, Chalivendra S, Logan D, Mattoo A, Selvaraj G, Datla R (2012) Target of Rapamycin Signaling Regulates Metabolism, Growth, and Life Span in Arabidopsis The Plant Cell 24: 4850–4874

Siqueira JA, Hardoim P, Ferreira PCG, Nunes-Nesi A, Hemerly AS (2018) Unraveling Interfaces between Energy Metabolism and Cell Cycle in Plants. Trends in Plant Science 23: 731–747

Solmonson A, DeBerardinis RJ (2018) Lipoic acid metabolism and mitochondrial redox regulation. J Biol Chem 293: 7522–7530

Somerville CR, Ogren WL (1982) Genetic modification of photorespiration. Trends in Biochemical Sciences 7: 171–174

Ströher E, Grassl J, Carrie C, Fenske R, Whelan J, Millar AH (2016) Glutaredoxin S15 Is Involved in Fe-S Cluster Transfer in Mitochondria Influencing Lipoic Acid-Dependent Enzymes, Plant Growth, and Arsenic Tolerance in Arabidopsis. Plant Physiol 170: 1284–1299

Taylor NL, Heazlewood JL, Day DA, Millar AH (2004) Lipoic acid-dependent oxidative catabolism of alpha-keto acids in mitochondria provides evidence for branched-chain amino acid catabolism in Arabidopsis. Plant Physiol 134: 838–848

Venne AS, Solari FA, Faden F, Parei T, Dissmeyer N, Zahedi RP (2015) An improved workflow for quantitative N-terminal charge-based fractional diagonal chromatography (ChaFRADIC) to study proteolytic events in Arabidopsis thaliana. Proteomics 15: 2458–2469

Vitha S, Beneš K, Phillips JP, Gartland KMA (1995) Histochemical GUS Analysis. *In* KMA Gartland, MR Davey, eds, Agrobacterium Protocols. Springer New York, Totowa, NJ, pp 185-193

Wiklund S, Johansson E, Sjöström L, Mellerowicz EJ, Edlund U, Shockcor JP, Goories J, Moritz T, Trygg J (2008) Visualization of GC/TOF-MS-Based Metabolomics Data for Identification of Biochemically Interesting Compounds Using OPLS Class Models. Analytical Chemistry 80: 115–122

Zhang Y, Primavesi LF, Jhurreea D, Andralojc PJ, Mitchell RA, Powers SJ, Schluepmann H, Delatte T, Wingler A, Paul MJ (2009) Inhibition of SNF1-related protein kinase1 activity and regulation of metabolic pathways by trehalose-6-phosphate. Plant Physiol 149: 1860-1871

