## Supplemental Figures for "GCS-H2 is essential for growth as it acts as the main relay for mitochondrial lipoylation in heterotrophic tissues of *Arabidopsis thaliana*"

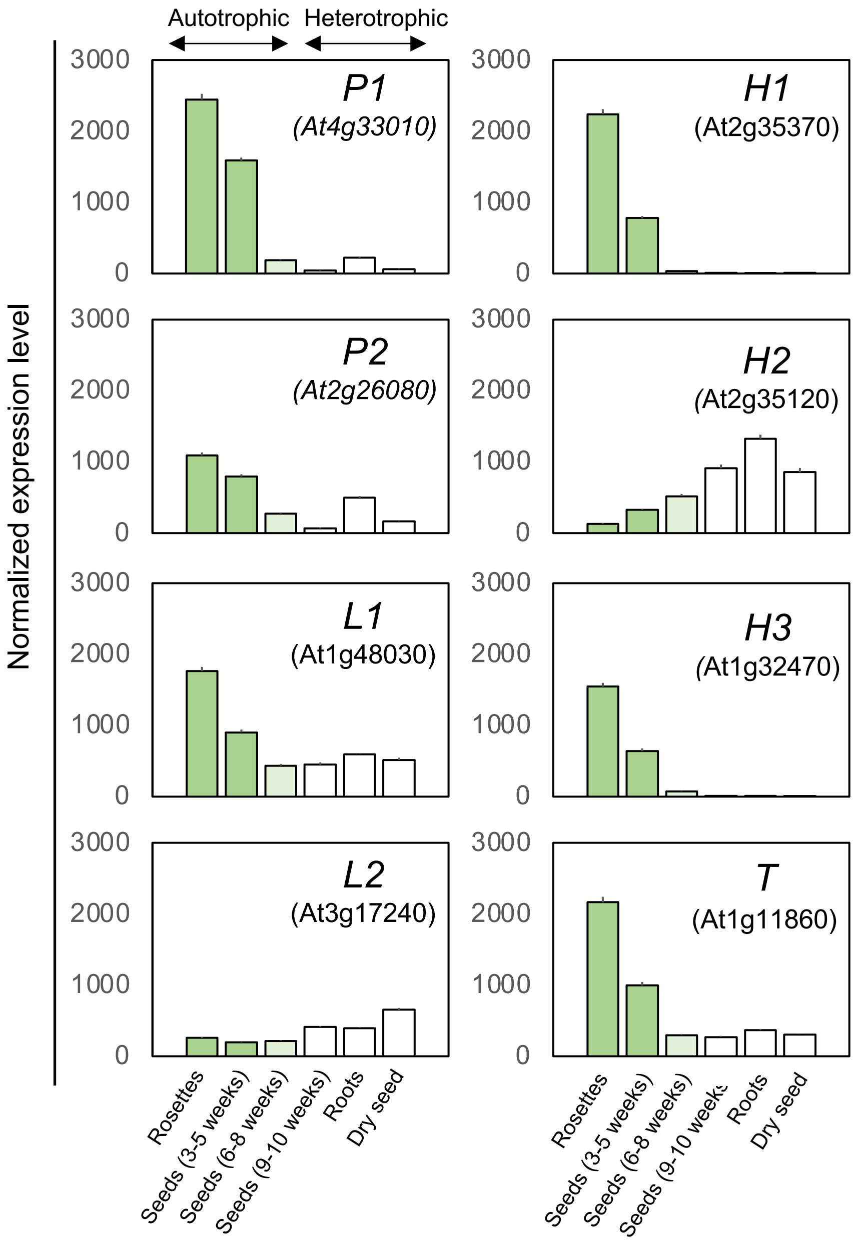


**Supplemental Figure 1.** Normalized transcript expression levels of genes encoding all isoforms of the four subunits of the Glycine Cleavage System in several autotrophic and heterotrophic Arabidopsis organs. Expression data were retrieved from AtGenExpress Consortium (Arabidopsis eFP Browser; http://bar.utoronto.ca/).

Begränsad delning

**a**

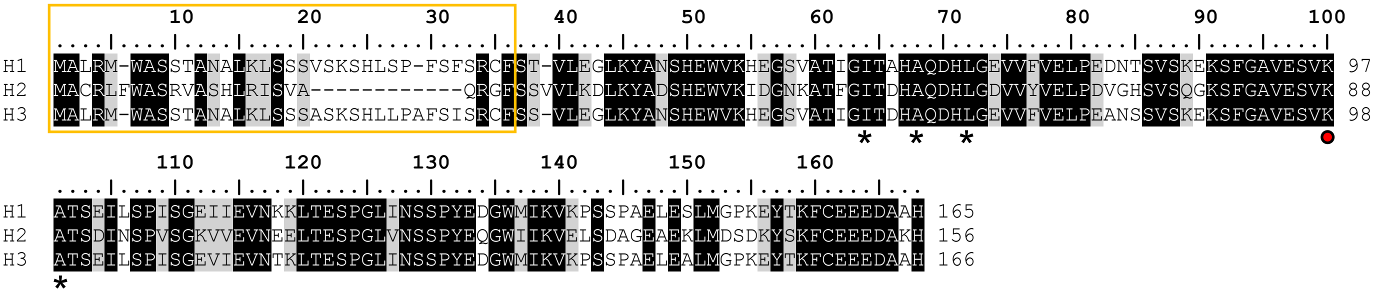


**b**

| | H1 | H2 |
|---|---|---|
| H2 | 67% | - |
| H3 | 95% | 69% |

**Supplemental Figure 2. Amino acid sequence analysis of Arabidopsis H proteins. (a)** Protein sequence alignment. Mitochondrial targeting sequences predicted by TargetP2.0 server are framed in orange. Conserved lipoyl-lysine is indicated by a red dot, whereas hydrophobic residues making contacts with the lipoamide chain (Cohen-Added et al., 1995) are indicated by an asterisk. Shaded areas indicate at least 90% amino acid identity (black) or similarity (grey). The alignment was obtained by using sequence alignment programs MEGA (MUSCLE algorithm) and BioEdit. **(b)** Identity percentages between H proteins. Sequence identity was calculated using ClustalW software (http://www.genome.jp/tools-bin/clustalw) from mature protein sequences (*i.e.* devoid of their mitochondrial targeting sequences).

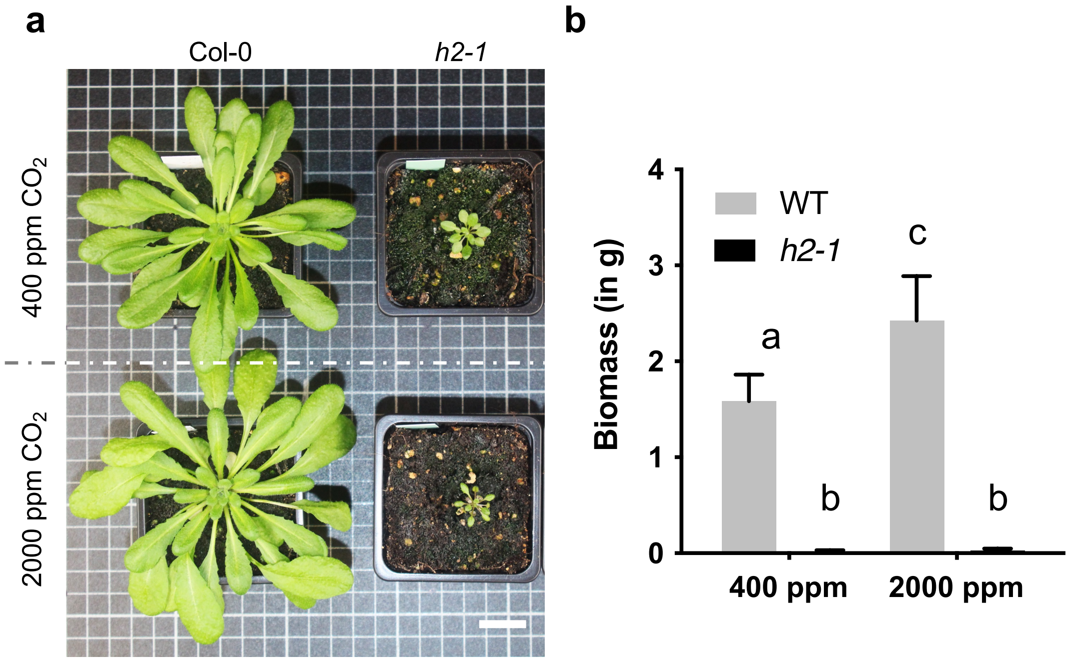


**Supplemental Figure 3. *h2* mutant phenotype under ambient air and high $CO_2$ conditions. (A)** Growth phenotypes of 7-week-old wild-type Col-0 (WT) and *h2-1* plants grown 5 weeks under short-day conditions at Ambient Air (400 ppm) or High $CO_2$ (2000 ppm) conditions. **(B)** corresponding fresh weight biomass in gram. Genotypes and conditions were marked as statistically different following a two-way ANOVA ($P<0.001$; $10<n<26$). Scale bar = 2 cm.

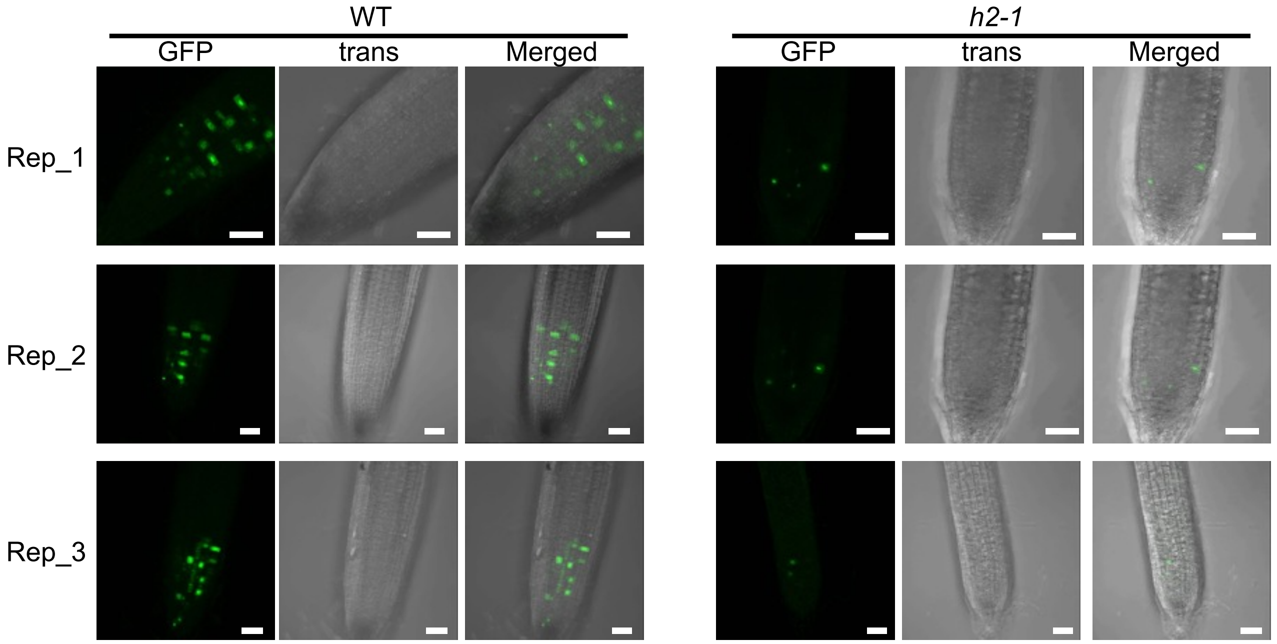


**Supplemental Figure 4.** Confocal images replicates for quantification of *pCycB1-eGFP* signals in root tips of *h2-1* and WT plants transformed with pcycB1-eGFP (pSL36 backbone, with Kanamycin changed to Hygromycin). Seedlings were grown on ½ MS + 0.1% sucrose during 8 days. Scale bars = 40 µm. Confocal microscope LSM780, gain (750); eGFP (488 nm/495-535 nm).

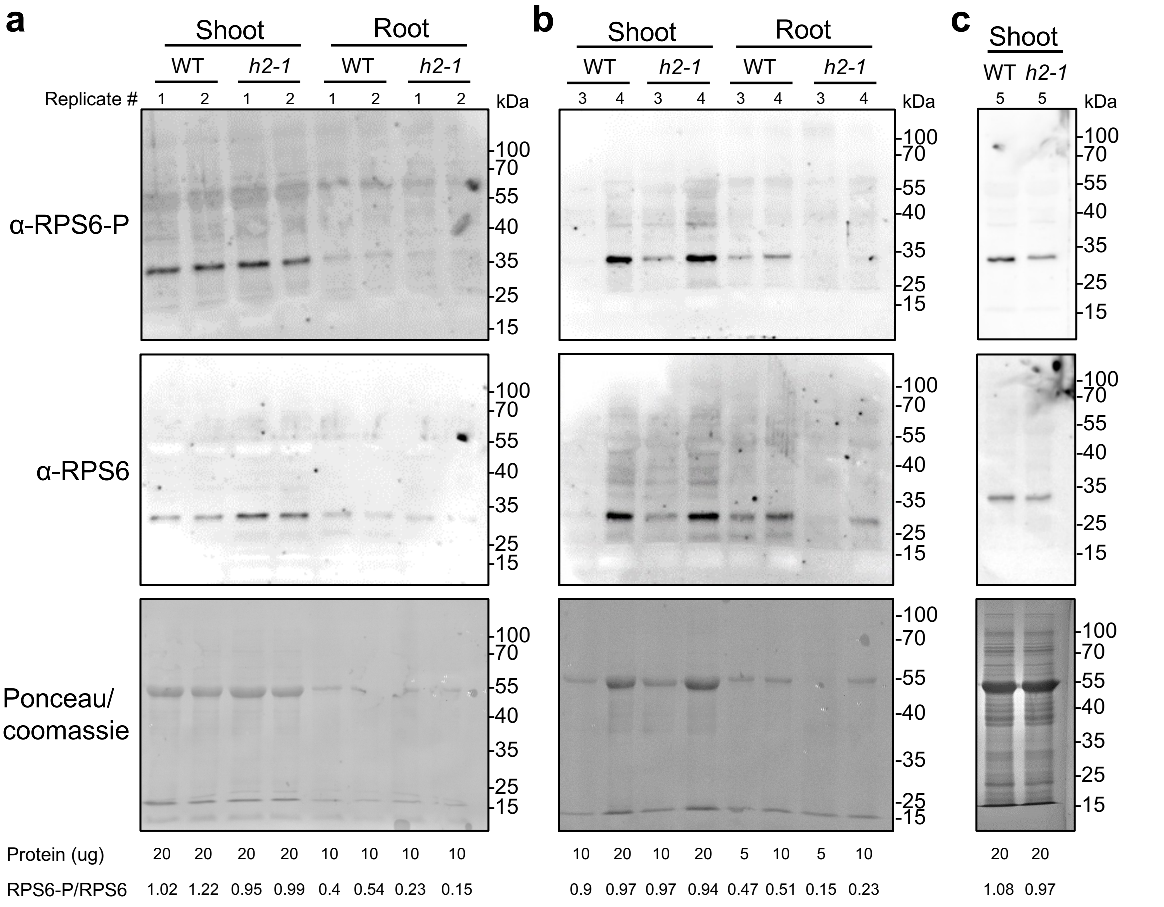


**Supplemental Figure 5. Gel replicates for RPS6-P/RPS6 for *h2-1* and WT plants.** Whole immunoblot gels and replicates reporting the phosphorylation state of RPS6, target of TOR, on 10 day-old shoots (n=5) and roots (n=4). The a, b and c panels show immunoblots (a: root/shoot replicates 1-2; b: root/shoot replicates 3-4; c: shoot replicates 5) and controls (a and b: ponceau; c: coomassie) from independent biological replicates.

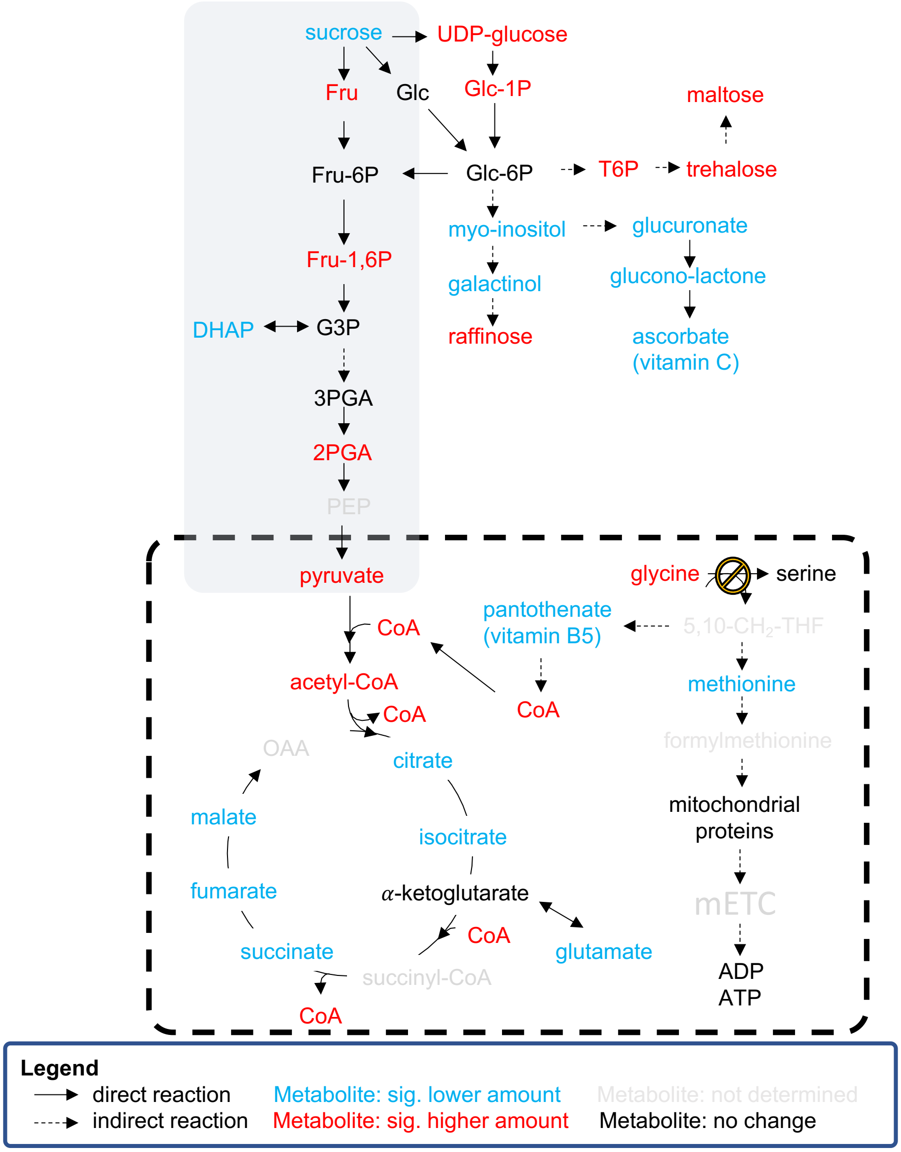


**Supplemental Figure 6.** Metabolite abundance in the primary carbon metabolism of roots in *h2-1* mutant as compared to WT. Red: metabolites found in a significantly higher abundance, blue: in a significantly lower abundance in *h2-1*.

**a**

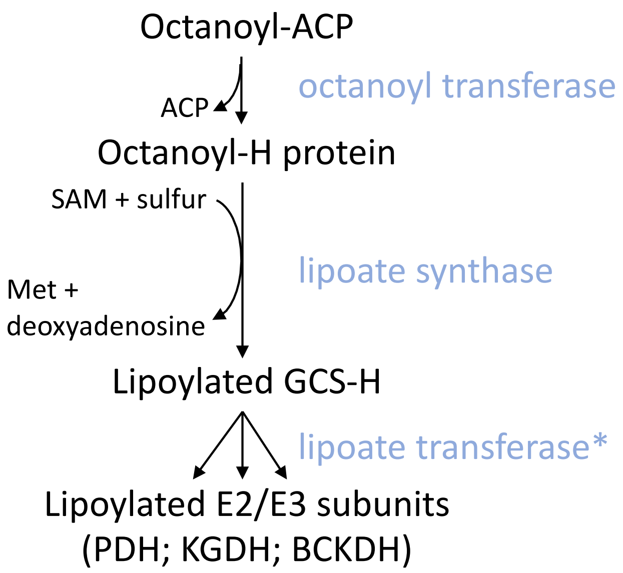


**b**

| | octanoyl transferase | lipoate synthase | lipoate transferase* |
|---|---|---|---|
| *E. coli* | LipB | LipA | - |
| *B. subtilis* | LipM | LipA | LipL |
| *S. cerevisiae* | Lip2 | Lip5 | Lip3 |
| *H. sapiens* | LipT2 | LiAS | LipT1 |
| *A. thaliana* | Lip2 | Lip1 | - |

**Supplemental Figure 7. (a)** Consensus *de novo* pathway for protein lipoylation in yeast (*S. cerevisiae*) and human (*H. sapiens*). **(b)** Nomenclature correspondence for enzymes involved in lipoic acid synthesis and transfer. * Lipoate transferase, also known as lipoyl amidotransferase, has not been reported neither in *E. coli* not in *A. thaliana*.
