## Supplementary material for "GCS-H2 is essential for growth as it acts as the main relay for mitochondrial lipoylation in heterotrophic tissues of *Arabidopsis thaliana*": Table S1_Primers

**Supplemental Table 1. List of primers used in this study.**

| Primer name | Primer sequence | Observations |
|---|---|---|
| SacI_pSL36 for | AGAGAGCTCGGGAAGTTTTCACTT | SacI + pSL36 (2182 bp 5' of the kanamycin ATG) |
| XbaI_pSL36 rev | ATCTAGAGTCGACCGGCATG | XbaI + pSL36 (687 bp 3' of the kanamycin STOP) |
| pSL36_Hygro for | AGGGCAAAGGAATAGGCGGGACTCTGGGGT | 15 nt Hygromycin pUBDEST + 15 nt pSL36 in 3' of Kanamycin |
| pSL36_Hygro rev | TTCAGGCTTTTTCATGCGAAACGATCCAGA | 15 nt Hygromycin pUBDEST + 15 nt pSL36 in 5' of Kanamycin |
| Hygro_pSL36 for | TCTGGATCGTTTCGCATGAAAAAGCCTGAA | 15 nt Hygromycin pUBDEST + 15 nt pSL36 in 5' of Kanamycin |
| Hygro_pSL36 rev | ACCCCAGAGTCCCGCCTATTCCTTTGCCCT | 15 nt Hygromycin pUBDEST + 15 nt pSL36 in 3' of Kanamycin |
| pSL36_Gen for | ATCGTTCAAATCCGTGAAGC | Genotyping and sequencing primers |
| pSL36_Gen rev | GCTTCACGGATTTGAACGAT | Genotyping and sequencing primers |
| Ampi_Gen for | GCTATGTGGCGCGGTATTAT | Genotyping and sequencing primers |
| Ampi_Gen rev | ATAATACCGCGCCACATAGC | Genotyping and sequencing primers |
| Hygro for | GTGCTTGACATTGGGGAGTT | Genotyping and sequencing primers |
| Hygro rev | GATGTTGGCGACCTCGTATT | Genotyping and sequencing primers |
| pcycB1 for | GAACAACGAACCGGAAAAGA | Genotyping and sequencing primers |
| AtH1 for | GGGGACAAGTTTGTACAAAAAAGCAGGCTTCATGGCACTAAGAATGTGGGCTTCT | Cloning in pUb-DEST |
| AtH1 rev | GGGGACCACTTTGTACAAGAAAGCTGGGTCCTAGTGAGCAGCATCTTCCTCCTC | Cloning in pUb-DEST |
| AtH2 promo for | GGGGACAAGTTTGTACAAAAAAGCAGGCTTCTTATCTCGCCAGGGATGTCCA | Cloning in pDONR207, pGWB1, pGWB4 (1000 nt upstream of the initial codon) |
| AtH2 rev | GGGGACCACTTTGTACAAGAAAGCTGGGTCGTGCTTGGCGTCTTCTTCTTC | Cloning in pDONR207, pGWB4 (without STOP codon) |
| AtH2 rev2 | GGGGACCACTTTGTACAAGAAAGCTGGGTCCTAGACCTACGGAGGTAAGGT | Cloning in pDONR207, pGWB1 (500 nt downstream of the stop codon) |
| AtH3 for | GGGGACAAGTTTGTACAAAAAAGCAGGCTTCATGGCACTGAGAATGTGGGCTTCC | Cloning in pUb-DEST |
| AtH3 rev | GGGGACCACTTTGTACAAGAAAGCTGGGTCCTAGTGAGCAGCGTCTTCTTCTTC | Cloning in pUb-DEST |
| AtH2 for | GCAAAAAAAAAAGCAAATCC | RT-PCR |
| AtH2 rev | GGTAACTCCACATAGACCAC | RT-PCR |
| AtACTIN-2 for | | RT-PCR |
| AtACTIN-2 rev | | RT-PCR |
| SALKseq_120199.2 for | TGCCAACAACAACAACAAAAG | *h2-1* genotyping |
| SALKseq_120199.2 rev | AGACGTTGTGACATGGACCTC | *h2-1* genotyping |
| SAIL_1152_G07 for | TTAGGGCAGTTCACGAAAATG | *h2-2* genotyping |
| SAIL_1152_G07 rev | TAAACAACATGTTTCAGGCCC | *h2-2* genotyping |
| SALK LBb1.3 | ATTTTGCCGATTTCGGAAC | *h2-1* genotyping |
| SAIL LB1 | GCCTTTTCAGAAATGGATAAATAGCCTTGCTTCC | *h2-2* genotyping |
